# The Pvc15 D2-Pnf SP Interaction Mediates AAA ATPase Activity, Payload Stability, and Translocation into the PVC Tube Lumen

**DOI:** 10.1101/2023.08.15.553202

**Authors:** Rhys Evans, Nicholas R. Waterfield

## Abstract

The *Photorhabdus* virulence cassette (PVC) is an elegant, multi-protein, contractile nanostructure which injects functionally bioactive polypeptides into the eukaryotic cytosol. Signal peptides (SPs) are N-terminal amino acid motifs from native ‘payload’ proteins which can also associate wide range of heterologous proteins to the PVC tube lumen. In addition, Pvc15 is a classic AAA ATPase encoded within the PVC operon which encodes an N-domain and tandem AAA domains D1 and D2. This work finds that Pvc15’s ATPase activity is mediated by the E555 residue situated in the Walker B motif of the sole functional ATPase domain, D2. Pvc15 multimerisation may be required to translocate payload into the PVC tube lumen, which is facilitated by intact N and D1 domains. In addition, ATPase activity requires the presence of other PVC operon components which hints at a PVC-regulated mode of action for loading. Furthermore, D2 is a chaperone for the native Pnf SP, though C-terminal truncations of the Pnf SP confers Pvc15-independent stability to the payload. These findings provide insight into the nuanced roles of the Pvc15-SP interaction for payload-PVC association and the effects that SP length and type has on payload stability and PVC-loading ability.

## Introduction

The *Photorhabdus* virulence cassette (PVC) is one classic example of a class of bacterial bioactive protein delivery devices collectively termed extracellular contractile injection systems, or eCIS (Yang *et al*., 2006; Hapeshi and Waterfield, 2017; Chen *et al*., 2019). Devices of this nature are often large (>10MDa) multiprotein phage-like nanosyringes consisting of a rigid, hollow tube whose distal spike is used to puncture into the cell membrane of the target host cell, after contraction of the surrounding interlinked sheath proteins, to deliver a toxic effector ‘cargo’ or ‘payload’ (Kube and Wendler, 2015; Desfosses *et al*., 2019; Jiang *et al*., 2019). *Photorhabdus* species encode 5-6 PVC operons in their genomes, each differing only slightly in the conservation of the PVC core structural components (Yang *et al*., 2006). Operons differ more greatly by the effector cargoes that are encoded up to ∼6kb downstream of their respective PVC operons and which give namesake to the operon itself. Cargo loading is mediated, in part, by the presence of a 50-70 amino acid N-terminal “signal peptide” (SP) which is effective for loading even large heterologous proteins such as β-lactamase, mRFP, and *Renilla* luciferase into the 150-200nm-long tube lumen (Jiang *et al*., 2022; Wang *et al*., 2021).

The modular nature of PVC payloads to be associated with the PVC offers an exciting potential for these devices to be bioengineered for the delivery of therapeutic proteins into the cytosol (reviewed by Lee *et al*., 2019). For example, the distal C-terminal domain of Pvc13 tail fibres can be replaced with epitope tags or adenovirus binding domains to retarget wild-type PVCs to various cell types (Kreitz *et al*., 2023).

How the PVC-specific SPs target the cargo to the PVC for loading remains a mystery. Pvc15, an <u>A</u>TPase <u>a</u>ssociated with diverse cellular <u>a</u>ctivities (a AAA+ ATPase), is required for cargo loading and for the unfolding of SP-associated mRFP (Jiang *et al*., 2022). In addition, the *Serratia entomophila* homologue, anti-feeding prophage (Afp) are only produced when co-expressed with Afp15, whereas Δ*pvc15* PVCs are intact but lack their associated cargo (Bhardwaj, Mitra and Hurst, 2021).

To understand the relevance of SPs to eCIS loading, this work presents a comprehensive analysis of eCIS effector diversity in lineage Ia and Ib operons, and protein prediction is used to identify the ubiquity of SPs in these ORFs. Protein structure prediction is also used to infer the function of Pvc15 domains in its monomeric and potential hexameric assemblies (Snider, Thibault and Houry, 2008). In addition, analysis of Western blot band intensities is used to infer the effects of SP type (i.e., Pnf50 and Cif50) and length on cargo stability and PVC loading when co-expressed with Pvc15 in *E. coli*. Finally, Pvc15 mutants are constructed and expressed *in trans* to the PVC*pnf*Δ*pvc15* operon where the ATPase activity is then characterised to infer the importance of the identified domains and residues to Pvc15’s various roles.

## Results and Discussion

### eCIS lineage analysis reveals diverse putative effector functionality

To assess the diversity of functions of effectors at eCIS operons, the eCIS database (dbeCIS; http://www.mgc.ac.cn/dbeCIS/) was used to identify lineage Ia and lineage Ib eCIS (Sarris *et al*., 2014; Chen *et al*., 2019). It was hypothesised that effectors associated with eCIS would be encoded up to 6kb downstream of the final ‘core’ gene in the operon: usually the Afp16 ‘cap’ equivalent gene. By appending the sequence information to a custom database, the function of each ORF was predicted by sequence and domain architecture using BLASTp and HHPred, respectively. Occurrences were then counted using regular expressions. The distributions of predicted functions in lineage Ia and Ib are shown in FIG. 1a and b, respectively.

**FIG. 1.**
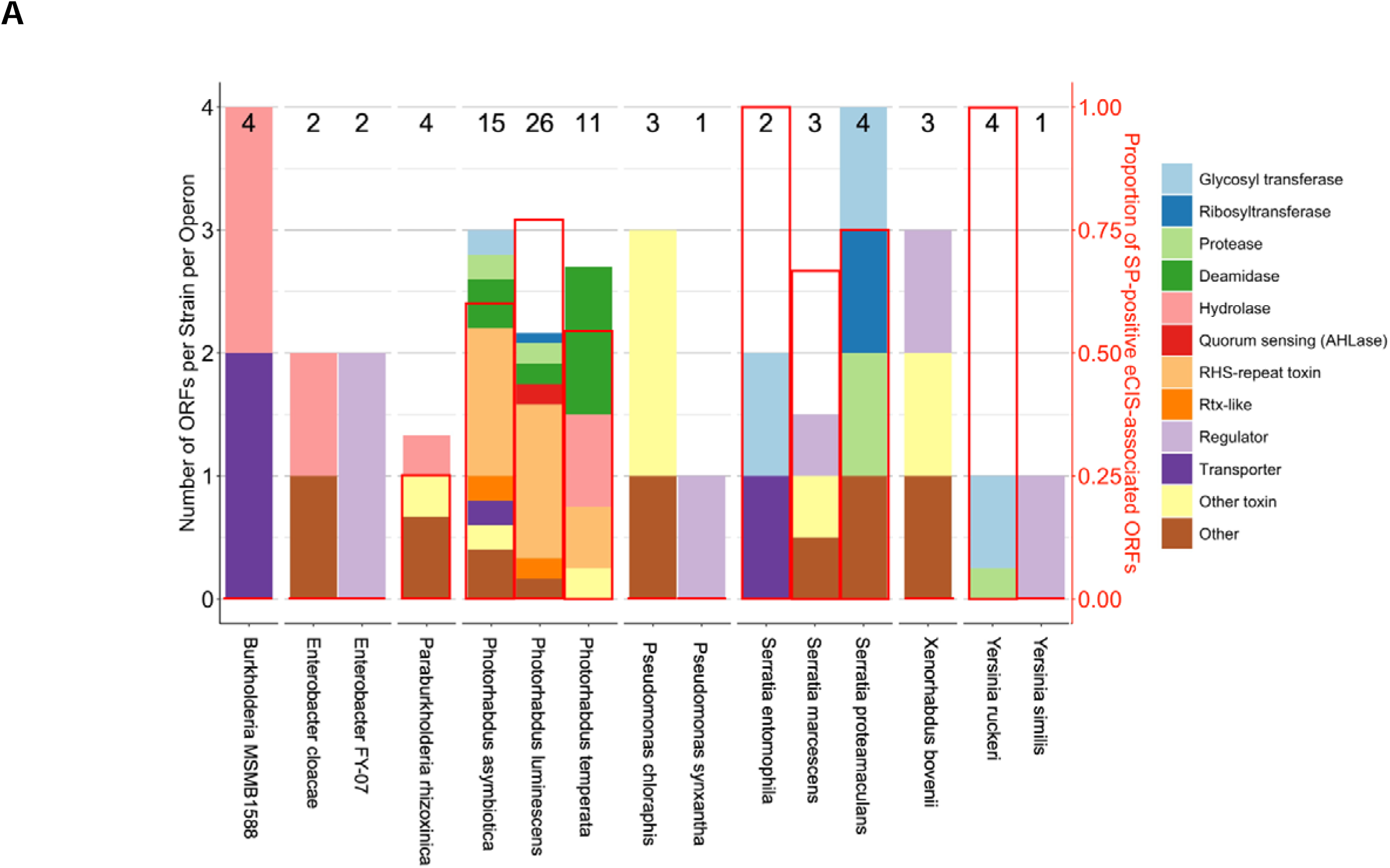
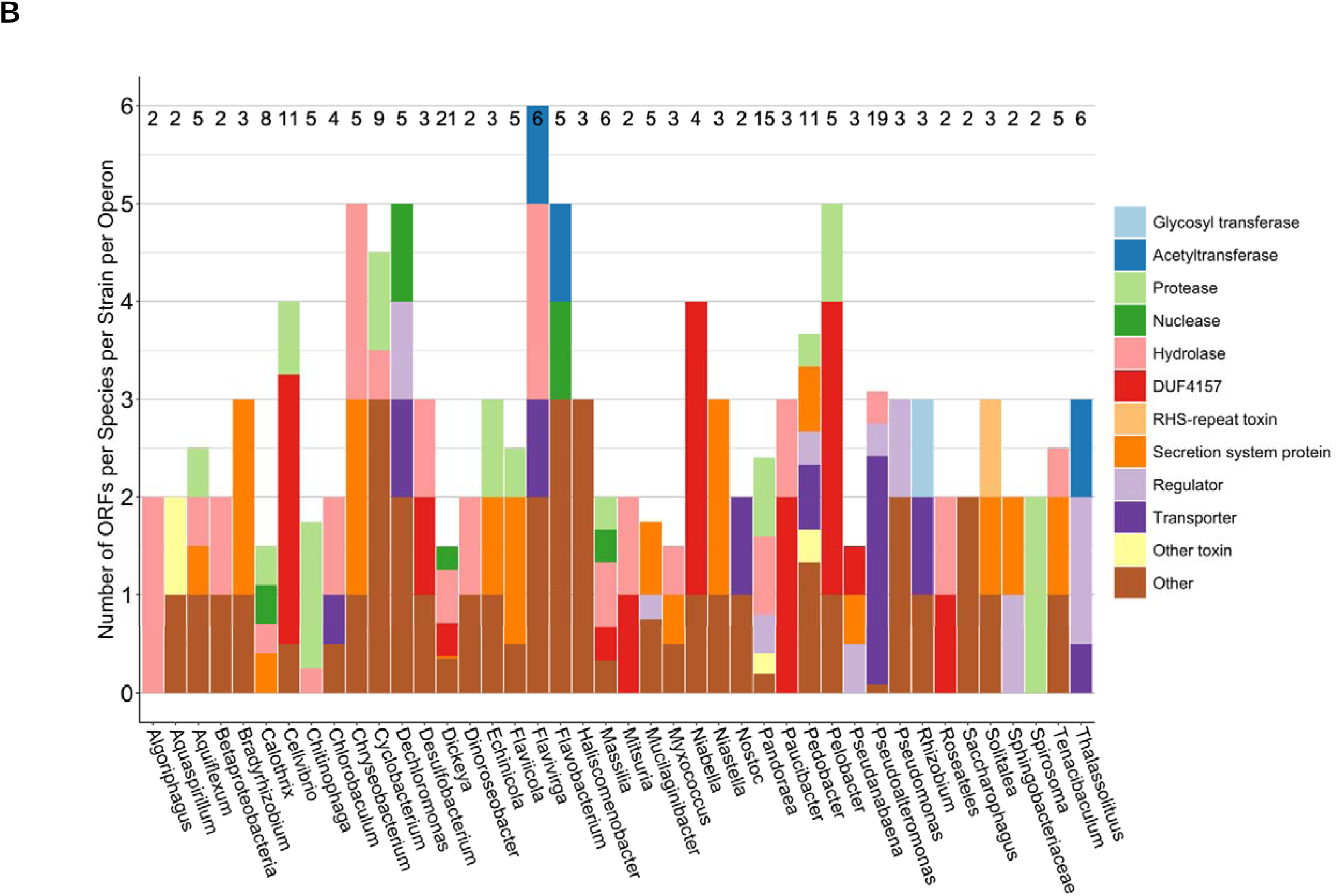
Taxonomic distributions of putative functions of lineage Ia and Ib eCIS-downstream ORFs. The dbeCIS database was used to identify structural operons of extracellular contractile injection system (eCIS) in lineage Ia **(A)** and Ib **(B)**, as defined by Chen *et al*., 2019. Open reading frames (ORFs) up to 6kb downstream of the final structural gene in the operon were appended to a database where BLASTp and HHPred were used to identify sequence homology and most likely function. Regular expressions were used to count how many times keywords appeared using R scripts. ORFs which did not fit any of the assigned keywords were classed as either ‘Other Toxin’ or ‘Other’ depending on additional information found for that entry. Numbers presented at the top of each bar represent the total number of ORFs used to generate the stacked bar. Functions were chosen based on the most commonly predicted functions in the database. AlphaFold2 predictions of each amino acid sequence was done to observe whether low confidence (i.e., highly disordered) N-terminal signal peptides (SPs) were present. The abundance of these sequences is displayed on the right axis with bars shown as red borders. In **(B)**, ‘Protease’ and ‘Nuclease’ are treated as subsets of ‘Hydrolase’ and are preferred to the more ambiguous ‘Hydrolase’ term. Genera encoding only one eCIS operon with one or less downstream ORFs were excluded from analysis: lineage Ia: *Chania* and *Shewanella*; lineage Ib: *Gynuella*, *Minicystis*, *Sorangium*, *Moorea*, *Geobacter*, *Dyadobacter*, *Draconibacterium*, *Anabaena*, *Aulosira*, *Tolypothrix*, *Fremyella*, *Herbaspirillum*, *Tenderia*, and *Stigmatella*.

*Photorhabdus* eCIS-downstream ORFs are significantly more highly represented in their use of rearrangement hotspot (Rhs) repeat-containing proteins, particularly in *P. asymbiotica* and *P. luminescens*. eCIS-downstream ORFs of the exclusively insect-pathogenic *P. temperata*, meanwhile, were more commonly predicted to have deamidase and hydrolase activities. It should be noted that the dbeCIS database does not include certain strains of *Photorhabdus*: *P. asymbiotica* HIT and JUN European strains and the Australian strain of *P. asymbiotica* Kingscliff (Wilkinson *et al*., 2010).

The 6kb cut-off in this methodology reflects the approximate sizes of each ORF within an eCIS operon as well as the distances between them. In total, there are 88 and 240 ORFs encoded up to 6kb downstream of the final base of the final core gene in the operon in lineages Ia and Ib, respectively. Of note, 24 ORFs across 13 species in lineage Ib (10% of all ORFs in lineage Ib) had either sequence or domain architecture similarity to a highly conserved domain of unknown function (DUF), DUF4157: a previously-termed PVC-metallopeptidase, although its function has never been confirmed. It contains a Zn-binding motif, HExxH, typical of metallopeptidases; authors have speculated its function to be involved in toxin interactions such as for loading or export (Geller *et al*., 2021). This hypothesis may be reflected in the current work since every instance of DUF4157 was encoded adjacent to another eCIS-downstream ORF with the only exception being *Stigmatella aurantiaca* strain DW4/3-1.

In addition, tertiary structures of ORFs of lineage Ia putative eCIS effectors were predicted using AlphaFold2 (Jumper *et al*., 2021) which was shown to be an accurate predictor of effector structure (FIG. S2). Signal peptides (SPs) could be identified by low-confidence N-terminal extensions (FIG. S2A and S2B). ORFs encoding SPs were verified under a set of conditions: amino acids 1-50 possessing an average pLDDT < 40 and considerably lower than that of the rest of the protein. Remarkably, SPs were found in 55.4% of ORFs up to 5kb downstream of a lineage Ia eCIS operon (FIG. 1A). These SPs are localised almost entirely to the PVC loci in *Photorhabdus* and the Afp loci in *Serratia* and *Yersinia*; this implicates the disordered SP as an Afp-homologue-specific feature of known eCIS cargoes.

### Protein prediction enables analysis of potentially relevant residues and supersecondary structures in the Pvc15 AAA domains

AlphaFold was used to predict the structure of the Pvc15 encoded at the Pnf operon: Pvc15*_pnf_* (Evans *et al*., 2021; Jumper *et al*., 2021; Mirdita *et al*., 2021). According to HHPred, Pvc15 is a putative AAA+ domain-containing protein most homologous to the mycobacterial homologue of Cdc48/p97/VPC and orthologue to *M. tuberculosis* Rv0435c: Msm0858 from *Mycolibacterium smegmatis* (5E7P) (Unciuleac, Smith and Shuman, 2016). Like Msm0858, Pvc15 has an N-domain, spanning residues 1-205, as well as two tandem AAA domains, D1 (residues 206-434) and D2 (residues 435-698) (FIG. S2; S2A by residue, S2B by secondary structure, S2C by inferred domains according to alignment (bottom), and S2D by pLDDT).

A closer look at the structure indicates that the conserved Walker A and B motifs are present in each of D1 and D2 (FIG. 2A). At the centre of the ATPase domain is a classical β-α-β sandwich or Rossmannoid fold (Hanukoglu, 2015), a hallmark of AAA+ proteins and ordered as β5-β1-β4-β3-β2 (Miller and Enemark, 2016).

**FIG. 2.**
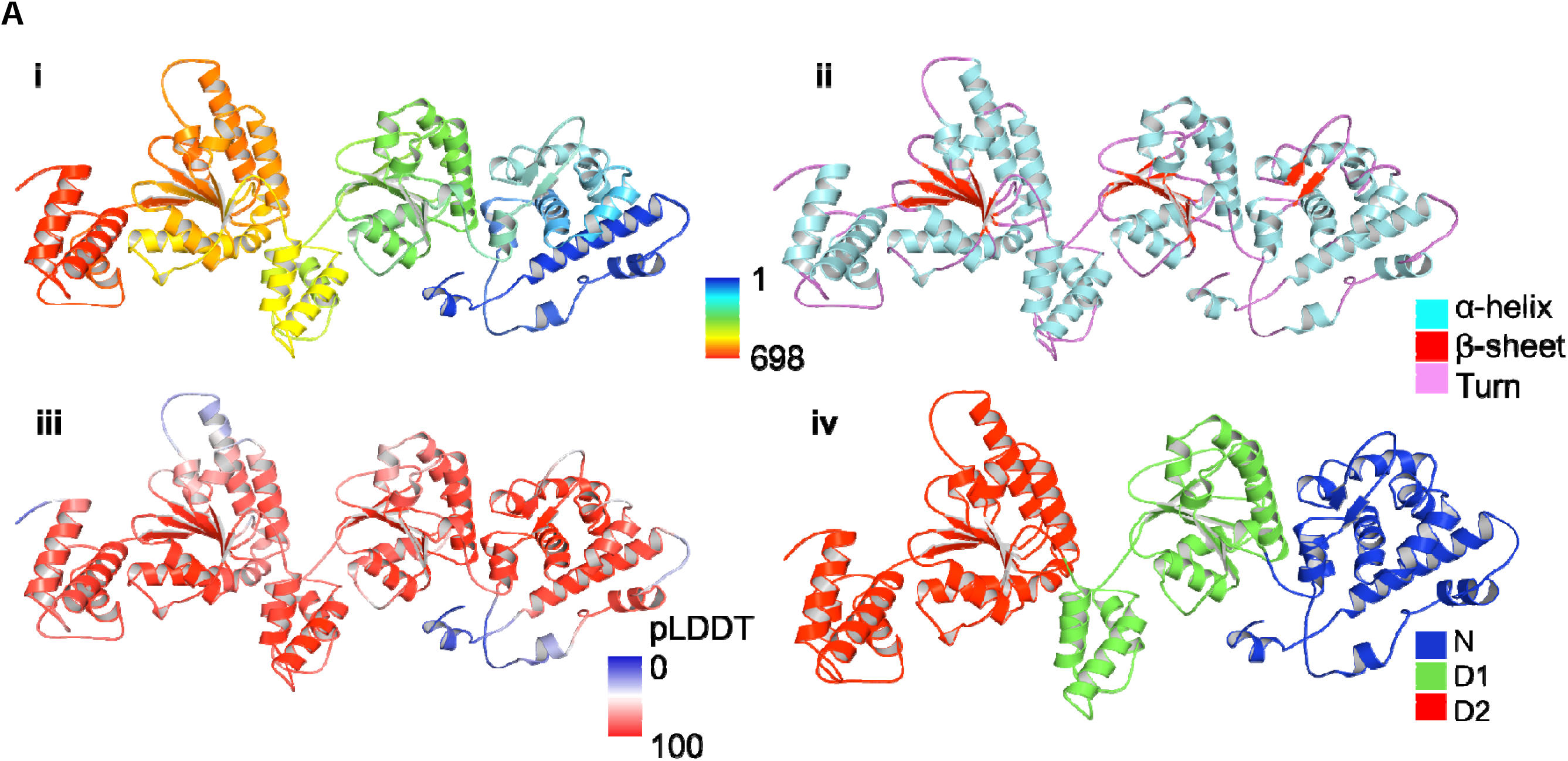
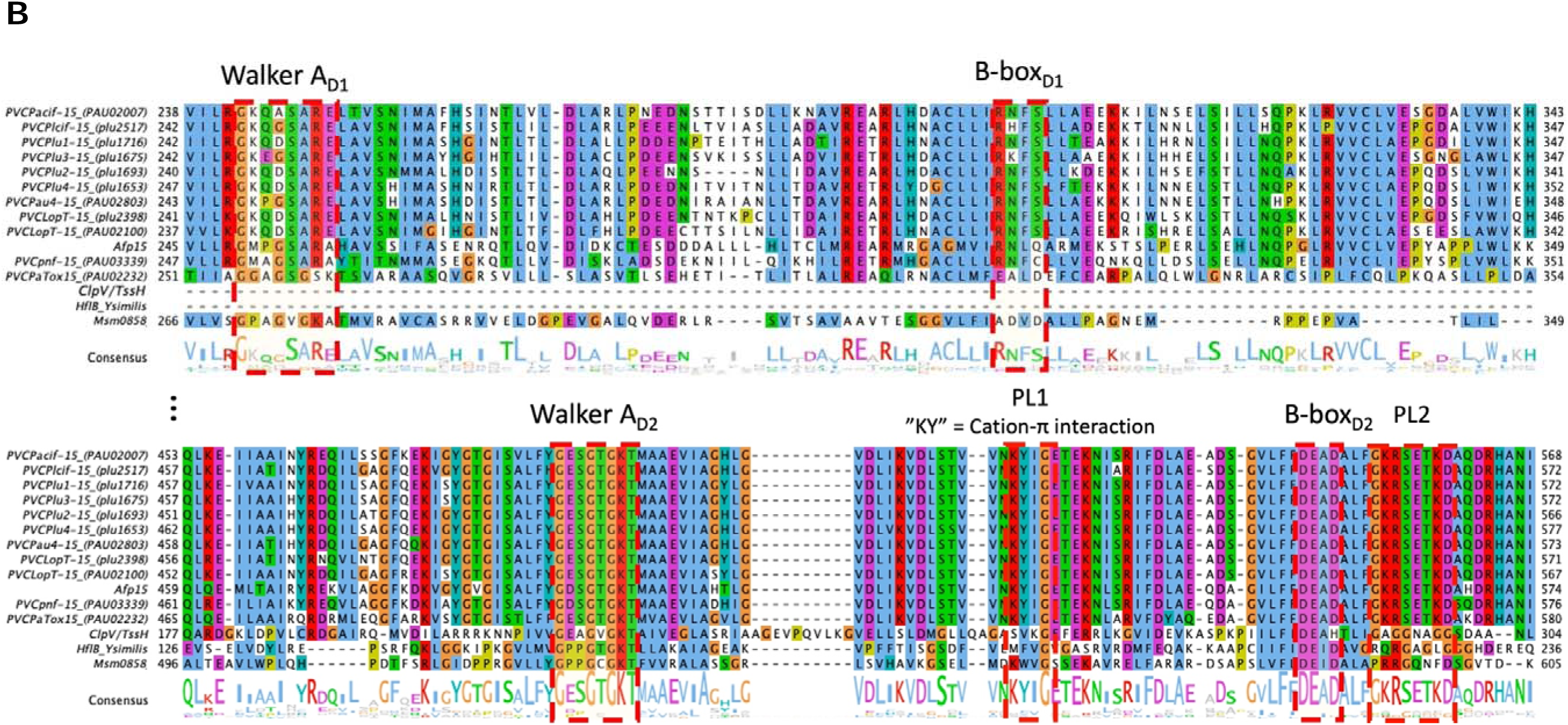
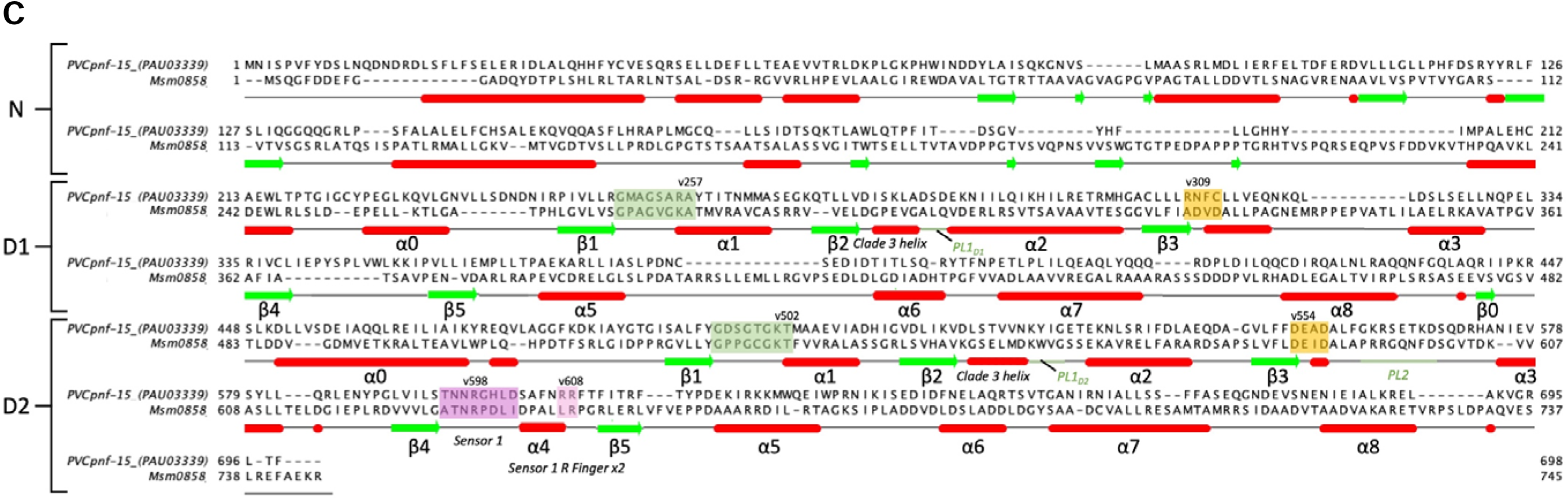
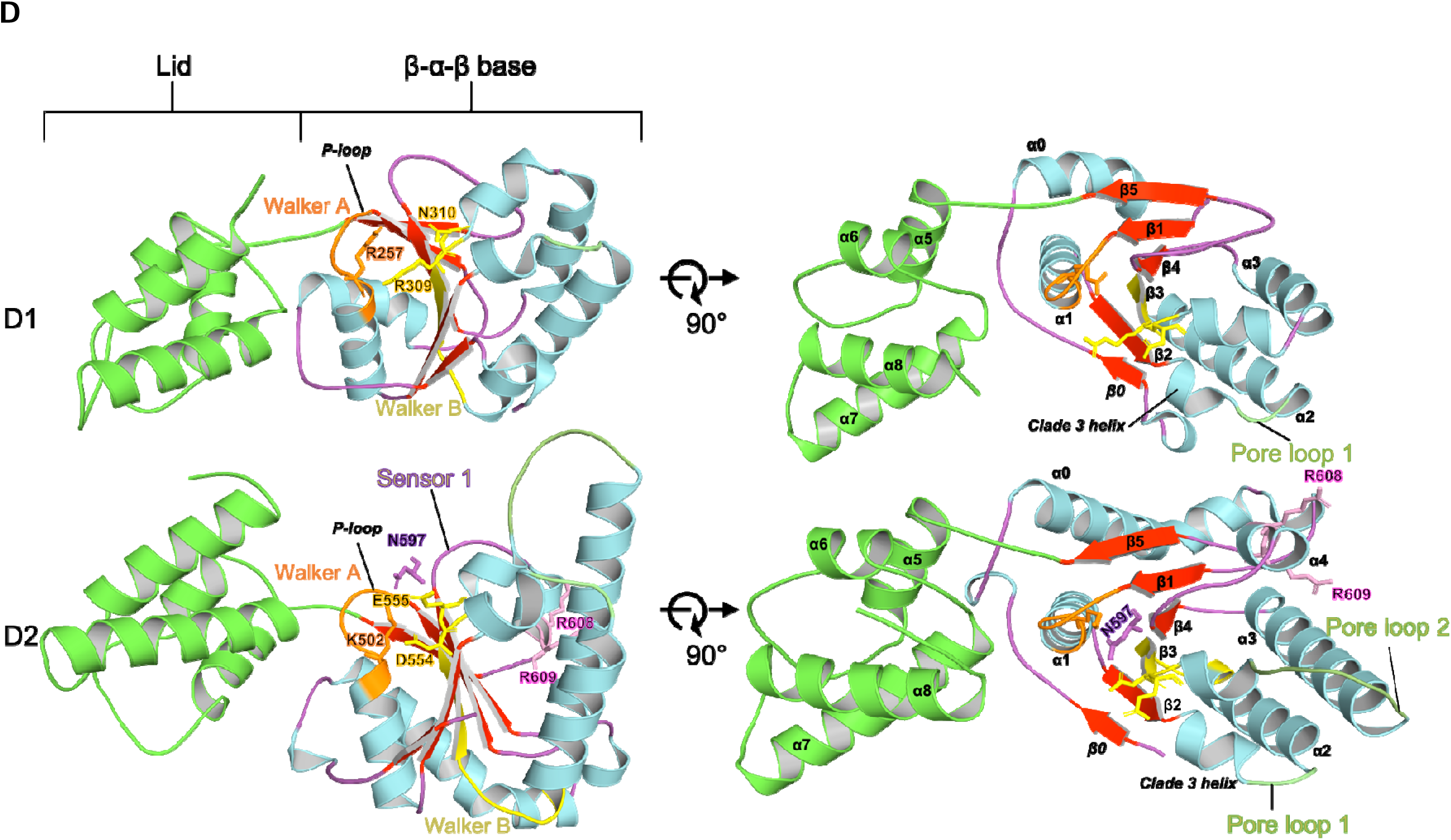
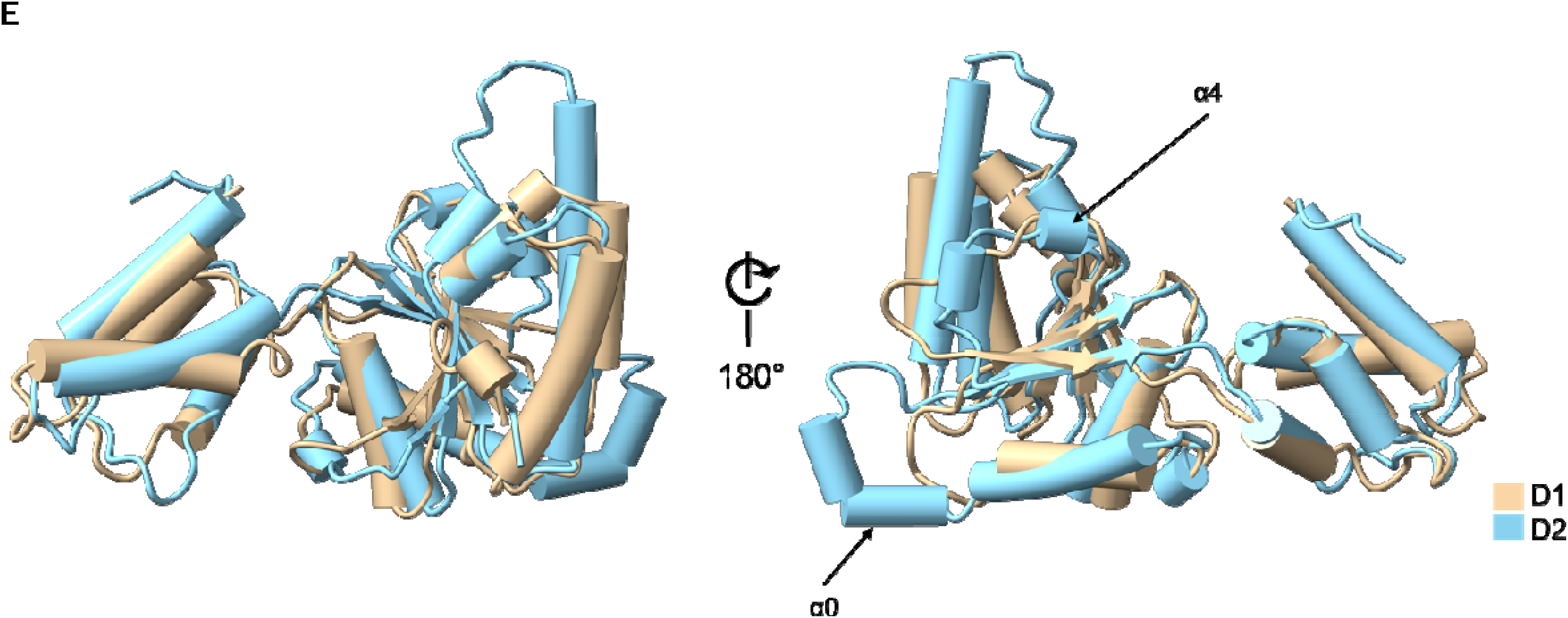
C-terminal ATPase domain in Pvc15 contains conserved features. **(A)** Walker A and B motifs are highly conserved ATP and GTP-binding sites and are found in the C-terminal region of Pvc15 homologues including Afp and ClpV/TssH. **(B)** Alignment of the α/β sandwich region of the C-terminal D2 domain of Pvc15. Notable motifs include the Walker A and B motifs which are highly conserved sequences of amino acids in AAA+ ATPases; these include the ‘nest’ residues Gly499-Lys502 in the Walker A motif, which coordinate ATP, and Asp554-Glu555 in the Walker B motif which coordinate a divalent cation to catalyse ATP hydrolysis. Other notable residues in the second region of homology (SRH) include the sensor 1 (S1) residue Asn597, the sensor 2 (S2) residue Arg659, and the more distant Arg-finger residues which coordinate neighbouring protomers in the quaternary structure: Arg608 and Arg609. **(C)** The Walker A motif “LRLR” nest with proposed coordination of phosphate as documented for the P-loop of Ras by Watson and Milner-White (2002). Phi bond angles are also shown; carbon atoms are shown in green, nitrogen in blue, and oxygen in red; Amber-relaxed side chains are depicted as thin sticks.

The ATPase domain supersecondary structure is typical of a clade 3 AAA+ ATPase (FIG. 2B; Miller and Enemark, 2016). In D2, the Walker A motif consists of a glycine-rich loop situated between β1 and α1. The “GTGK” motif forms a compound nest of the form “LRLR” which coordinates the β-phosphate of ATP or ADP within the concave of the NH bonds, as in the case with Ras (FIG. S4). Divalent cations such as Mg^2+^ may also coordinate the phosphate groups (Bugreev and Mazin, 2004). Lys502 is likely responsible for nucleotide binding; a role which can be abrogated by mutating to alanine in homologous structures (Hanson and Whiteheart, 2005).

Walker B motifs tend to be less well conserved but often feature a set of up to 4 hydrophobic amino acids followed by a ‘DE’; in Pvc15 these residues are part of a ‘DEAD’ motif, which is often found in proteins with helicase activity (FIG. 2C) (Watson and Milner-White, 2002; Cordin *et al*., 2004). The aspartate residue (D554) may coordinate a divalent cation such as Mg^2+^ whilst the glutamate residue (E555) is responsible for ATP hydrolysis. Mutation of the latter to a glutamine is sufficient to impair this process in other ATPases such as the Lon protease (Shin *et al*., 2020) and SecA (Kim *et al*., 2013). On the other hand, Jiang and colleagues (2022) identified the Walker A and B motifs of Pvc15*_pnf_*but apparently failed to abolish its unfolding activity upon SP-associated fluorescent proteins by point mutations, indicating that Pvc15 may have other roles outside of its ATPase activity.

Additionally, the second region of homology (SRH) is a defining characteristic of AAA+ ATPases (FIG. 2B; right): the sensor 1 motif is located in the loop between β4-α4; the sensor residue, Asn597 is likely responsible for orientation of the nucleophilic water used in hydrolysis by the Walker B motif. Finally, two arginine fingers, Arg608 and Arg609, are located in a position which may enable them to stabilise the negative charge of the γ phosphate during ATP hydrolysis within a neighbouring cleft (i.e., *in trans*) in the hexameric structure.

Overall, protein structure prediction indicates that Pvc15 is a type II class 3 ‘classical’ AAA+ ATPase. A search of the DALI server (Holm and Rosenström, 2010) using the N domain of Pvc15 indicates many hits for DNA-directed transcription factor subunits, the best result being RNA polymerase II transcription factor B from *S. cerevisiae* with a Z-score of 7.5 at 2.9Å r.m.s.d (PDB: 5OQJ) (Schilbach *et al*., 2017) and also includes the MepR transcriptional regulator from *S. aureus* with a Z-score of 6.8 and 3.5Å r.m.s.d over 90 aligned residues (PDB: 4LD5) (Birukou *et al*., 2013).

In the close type II orthologue from *M. smegmatis*, Msm0858, both the D1 and D2 domains are capable of ATP hydrolysis despite having deviant residues in the D1 domain Walker B box and sensor 1 residue (Unciuleac, Smith and Shuman, 2016). Meanwhile, the Cdc48/p97 homologue’s D1 domain is used for stabilising the hexamer and binding ATP whilst the D2 domain is the primary catalytic domain (Baek *et al*., 2013). All Pvc15 homologues have even more deviant residues than these homologues: Pvc15 lacks even an aspartate or glutamate residue in the Walker B box, the Walker A nest lysine is replaced with arginine, and it lacks any second region of homology (i.e., sensor 1 and R-fingers). Altogether, these data indicate that D1 is likely deficient for ATPase activity but is, nonetheless, required for oligomerisation of Pvc15 to form a functional ATPase hexameric complex.

### Pvc15 hexamer predictions suggest an asymmetrical arrangement with pore poops for substrate translocation

MoLPC was used to predict the most likely complete hexameric assembly of Pvc15 (FIG. S5) (Bryant *et al*., 2022). The resulting structure represents an asymmetrical structure with a pore diameter of 13.7Å at the D2 domains (FIG. S5A). N-terminal domains assume a more open conformation and each subunit neighbours the next at an angle of between 9° and 12°; this produces an open right-handed ‘spiral staircase’ arrangement. Two of the subunits at the ‘top’ and ‘bottom’ of the staircase are predicted to be adjacent at an angle of ∼46°.

Pore loops 1 and 2 spiral in accordance with the larger domains and represent two ‘faces’ capable of contact with the translocating substrate (FIG. S5B). Lys528 and Tyr529 form a cation-π interaction, which suggests a strong binding to substrate, whilst Arg657 may play a role in contacting negatively charged amino acids in the substrate. This arrangement is similar to that of Vps4 (PDB: 6BMF) though the environment surrounding the aromatic residue can be specific for the ATPase’s intended function; the grip would be looser when neighbouring glycine (e.g. ClpB D2 (PDB: 6OAY)) or made more hydrophobic when neighbouring aliphatic residues such as valine (e.g. YME1 (PDB: 6AZ0)) (Puchades, Sandate and Lander, 2020).

Moreover, Arg608 and Arg609 are predicted to be located in a locale where they may contact the ATP-binding site *in trans* to communicate macromolecular changes during hydrolysis to the adjacent subunit (FIG. S5C).

### Different signal peptides have differential effects on cargo stability

Jiang and colleagues (2022) characterised Pvc15 as being necessary for loading of cargo proteins to the PVC. To establish whether this effect was due to cargo stabilisation or loading efficiency, it was investigated what how the SP affects cell lysate abundance.

To first ascertain the optimal time to measure protein expression after induction in NiCo21 cells, aliquots taken over time were analysed by Western blot and normalised by sum of lanes in a Coomassie stain. 3-4 h post-induction had the highest levels of protein expression (FIG. S6), so subsequent experiments on cargo stabilisation were conducted at 3 h post-induction with IPTG and arabinose to induce both the cargo and PVC*pnf* operon.

PVC*pnf* with either native Pnf or Pnf toxoid (C190A) were induced in the presence or absence of Pvc15 and normalised using the FLAG-tagged Pvc16 in Western blot analysis (FIG. 3A). Normalised band intensities on these Western blots showed a significant decrease in the stability of Pnf when Pvc15 was removed (t(4) = -5.55, p = 0.005 **), indicating that Pvc15 does act to increase protein abundance in the lysate. Surprisingly, however, truncation of the SP from its C-terminus made the cargo more stable than when it was encoded with the native SP, regardless of whether Pvc15 was absent (t(4) = 3.30, p = 0.030 *) or present (t(4) = 4.26, p-value = 0.013 *).

**FIG. 3.**
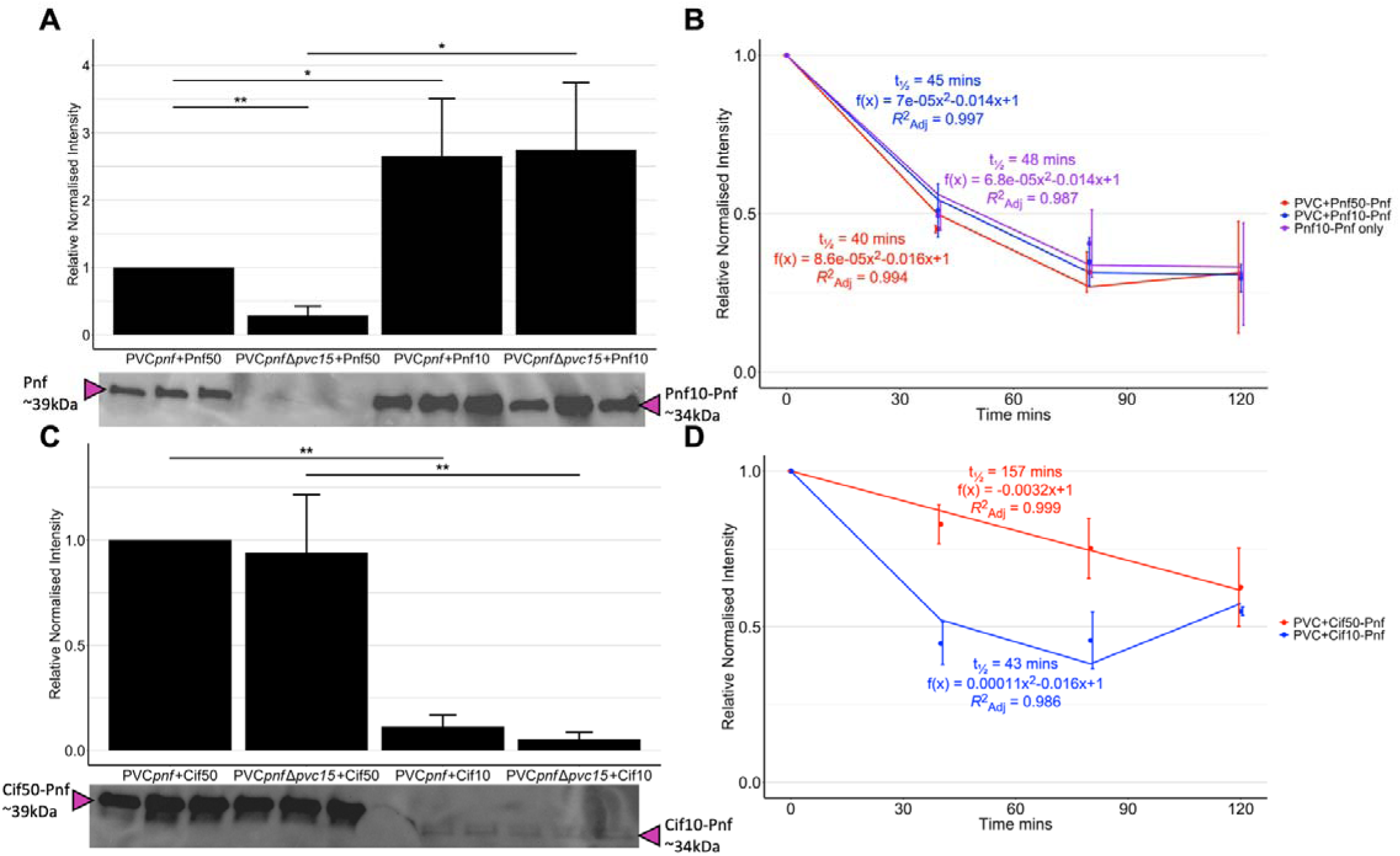
C-terminal truncations of signal peptides differentially affect the stability of the cargo. The abundance of Pnf-Myc was characterised using normalised band intensities from α-Myc + α-β-RNA polymerase (β-RNAP) primary antibodies. **(A)** *E. coli* BL21(DE3) cells were induced for 3 h at 25 °C with 0.2% arabinose and 0.5mM IPTG for PVC*pnf* and cargo induction, respectively. Error bars indicate standard deviation of 3 biological replicates (n = 3); statistical inference was done using a Student’s T-test. Deletion of Pvc15 from the PVC*pnf* operon results in a dramatic decrease in abundance of native Pnf50-Pnf-Myc from cell lysates (t(4) = -5.55, p = 0.005 **). However, C-terminal truncation of the Pnf SP to retain the 10 N-terminal amino acids (Pnf10) increases the abundance of the cargo by over 100% when co-expressed with the full PVC*pnf* operon (t(4) = 3.30, p = 0.030 *) as well as with the Pvc15-deleted operon (t(4) = 4.26, p = 0.013 *). Pnf10-Pnf-Myc abundance was unaffected by the presence or absence of the chaperone, implying a differential effect of the chaperone targeted towards the native 50 amino acid SP. **(B)** Cells induced for 3 h were centrifuged to remove IPTG and pelleted cells were resuspended in the same volume of LB + kanamycin at 20 μg/mL to halt both transcription and translation. Aliquots were then taken every 40 minutes. Non-linear regressions were predicted using the lm() function in R to compare coefficients and half-lives. C-terminal truncated Pnf10-Pnf decays at roughly the same rate as that of native Pnf regardless of its co-expression with the PVC (t_1/2_ = 40-48 mins). **(C)** For co-expression with the Cif SP (Cif50), truncation led to a decrease in abundance of the Pnf cargo, regardless of the presence of Pvc15 (with: t(3) = -11.28, p = 0.0013 **; without: t(4) = -5.51, p = 0.005 **). **(D)** The Cif SP truncation mutant (Cif10) displayed a significantly faster second-order degradation rate (t_1/2_ = 43 mins), compared to the native Cif SP (Cif50) which displayed a first-order degradation rate (t_1/2_ = 157 mins).

Measuring protein content after the addition of kanamycin at 20 μg/mL to block protein synthesis indicated that the effects on stability were due to differences in protein degradation rather than translation (FIG. 3C). Altogether, this indicates that Pnf alone is rapidly degraded and rescued only by the co-expression of Pvc15; of note, strains encoding Pvc15 displayed this effect regardless of whether Pvc15 was induced, indicating that Pvc15 may exert this chaperoning effect at very low cytosolic levels. Given this interpretation, Pvc15 is predicted to rescue roughly 30% of cytosolic Pnf from degradation (95% CI [15%, 43%]) when compared to the abundance of the Pvc15-independent SP, Pnf10.

Conversely, Pvc15 had seemingly no effect on the stability of Pnf encoded with either the native Cif50 SP or the C-terminal truncated Cif10, though the latter displayed a decrease to the stability of the cargo compared to the longer SP both with and without Pvc15 (t(3) = -11.28, p = 0.0013 **; t(4) = -5.51, p = 0.005 **) (FIG. 3D). This indicates that Pvc15 interacts differentially between the different PVC cargo SPs. In addition, the difference in band intensity observed between Pnf10 and Cif10 indicates that the 10 amino acid tip is, indeed, an important factor affecting the cargo’s stability. In contrast, the SP tip appears to be dispensable for Pvc15-dependent stabilisation since only Pnf50 displayed a differential effect with or without the chaperone.

### Signal peptides work with the Pvc15 ATPase for loading

If native Pnf can be stabilised by encoding shorter SPs, presumably making it a less desirable target for protein degradation machinery, then perhaps the longer SPs which are encoded naturally are preferred in the context of loading. Therefore, it was hypothesised that cargoes encoding C-terminal truncated SPs would be far less capable of being loaded into the PVC*pnf* chassis; this, then, would justify the bacterium having evolved longer, unstable signal peptides in its native cargoes to optimise loading whilst also encoding a nearby chaperone to mitigate the accompanying instability.

To compare loading efficiencies of the different SP-truncated cargoes, PVC*pnf* syringes were purified using immunoprecipitation (FIG. 4). There was a visual decrease in the relative level of Pnf in the final elution step of the α-Myc Western Blot (FIG. 4A). Overall, either truncation of the signal peptide or the deletion of Pvc15 resulted in a ∼50-75% decrease in the loading efficiency of Pnf (FIG. 4B). These results, therefore, confirm the hypothesis that shorter SPs, whilst conferring stability to the Pnf cargo, fail to recruit cargo to the PVC at comparable levels to that of the full-length SPs.

**FIG. 4.**
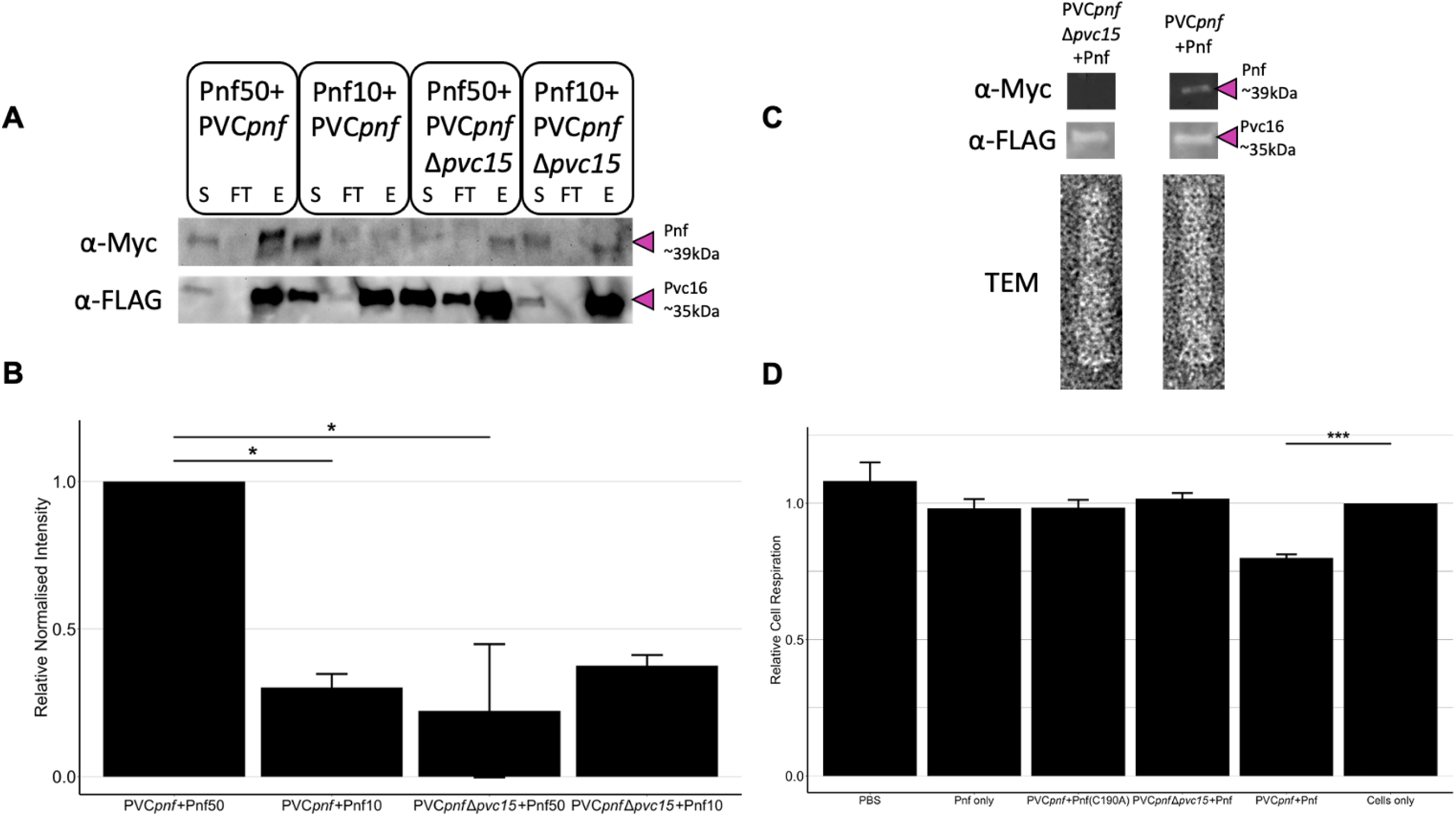
Full-length SPs and a Pvc15 chaperone are required for effective PVC*pnf* cargo association. pVTRa vectors expressing Myc-tagged Pnf with either the full-length (Pnf50) or the C-terminal truncated SP (Pnf10) were co-expressed with pBADPVC*pnf* or a mutant with deleted Pvc15 (PVC*pnf*Δ*pvc15*). PVC*pnf* nanosyringes were immunoprecipitated. **(A)** For each of the 4 cell strains, band intensities of the purification steps, supernatant (S), flow-through (FT), and elution (E), were compared by Western blot using α-Myc or α-FLAG primary antibodies. Band intensity for Pnf cargo in the elution step of the native system (Pnf50+Pvc15) was visually more pronounced. **(B)** α-Myc band intensity in **(A)** was normalised by the corresponding α-FLAG band. More than a 50% decrease in loading efficiency was observed for all mutants with either a C-terminal truncated SP (t(3) = -3.22, p-value = 0.049 *), or in the absence of Pvc15 (t(4) = -4.23, p-value = 0.013 *). Error bars show standard deviation with n = 2-3 reiterations of purified nanosyringes; statistical analysis was done with a two-sided Student’s T-test. **(C)** Western blot and TEM were used to verify the purification of intact PVC*pnf* after anion exchange. The PVC*pnf*Δ*pvc15* mutant failed to co-purify with cargo. **(D)** The effect on respiration of Pnf-loaded purified PVC*pnf*, as well as Ni-NTA-purified Pnf cargo, was tested on Raji B cells to measure resazurin reduction into the fluorescent resorufin. The native Pnf-loaded PVC*pnf* impeded the reduction of resazurin by the cells (t(6) = -9.95, p-value = 5.569e-05 ***) whilst other samples did not affect cell viability, including Pnf with a C190A mutation of the active site. Pnf co-expressed with the PVC*pnf*Δ*pvc15* operon similarly had no effect. Pnf-Myc alone was insufficient to affect resorufin fluorescence, indicating the importance of its PVC*pnf* vector for delivery.

After further purification of the PVCs, the absence of cargo was made even more apparent by Western Blot, though, as expected, the PVCs purified from the PVC*pnf*Δ*pvc15* operon were intact when imaged in transmission electron microscopy (TEM) (FIG. 4C).

As shown by recent data, PVCs can be artificially targeted to certain cell receptors to ensure optimal payload injection and cell type specificity whereby Pnf can be made to be cytotoxic to the target cells (Kreitz *et al*., 2023). However, some background binding to non-targeted cells is also possible. Pnf acts similarly to *E. coli* cytotoxic necrotizing factor, CNF, by deamidating a crucial residue in RhoA which subsequently causes the formation and maintenance of stress fibres (Buetow *et al*., 2001; Vlisidou *et al*., 2019); the cytoskeleton has been subsequently shown to interfere with mitochondrial activity via voltage-dependent anion channels (VDAC) and mitochondrial creatine kinase (MtCK) (Guzun *et al*., 2009). Therefore, the effects of Pnf-loaded purified PVCs (with native Pvc13 tail fibres) on eukaryotic cell respiration could be tested on Raji B cells by the addition of resazurin to measure fluorescence from the mitochondria-reduced form: resorufin (FIG. 4D). The optimal starting cell concentration for a 24 h time course was found to be 8 x 10^5^ cells/mL (FIG. S7).

These results also find that the SP does not require cleavage to be loaded into the PVC since purified samples resulted in Pnf molecular weights corresponding to uncleaved SPs (∼39kDa). Smaller bands observed in *E. coli* lysates (∼25kDa and ∼17kDa) are also present when expressing Pnf on its own without any other PVC components, indicating that these bands may be the result of Pnf degradation by the cell.

### Pvc15 domain D2 is an effective chaperone for the Pnf SP and is the sole functional ATPase domain

Guided by the bioinformatic analysis of Pvc15 structure, plasmids encoding either wild-type or mutant Pvc15 downstream of the same promoter as the Pnf payload, pVTRaDuet (FIG. S8), were made and transformed into *E. coli* BL21 (DE3) co-expressing PVC*pnf*Δ*pvc15*. Mutants include deletions of the N to D1 domains (Pvc15ΔN-D1), the D2 domain (Pvc15ΔD2), or abolishment of the catalytic E555 (Pvc15E555Q). Optimal Pnf abundance was measured by comparing Western Blot band intensities from lysates at 4 h post-induction which showed that deletion of domain D2 resulted in a depletion of Pnf payload, suggesting this domain is an effective chaperone (FIG. 5A). Unlike when Pvc15 is deleted, however, some Pnf can be observed which may indicate that domains N and/or D1 are also able to stabilise the payload somewhat.

**FIG. 5.**
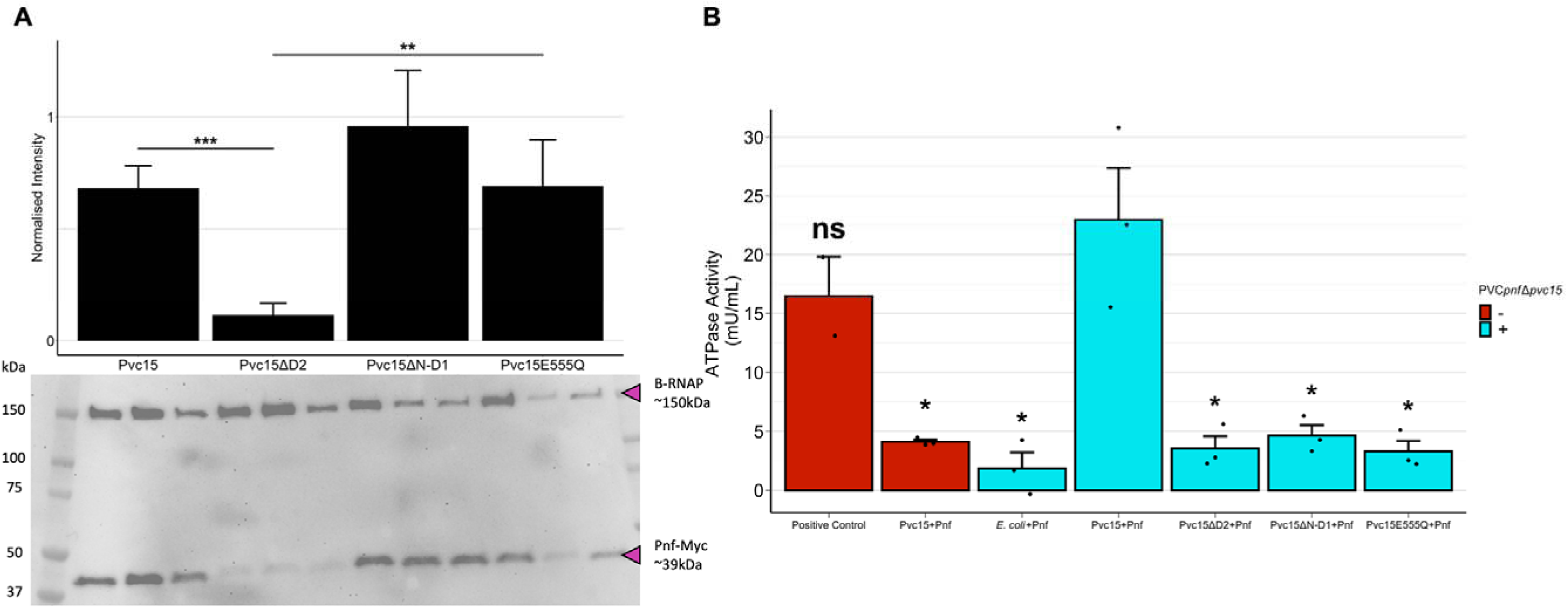
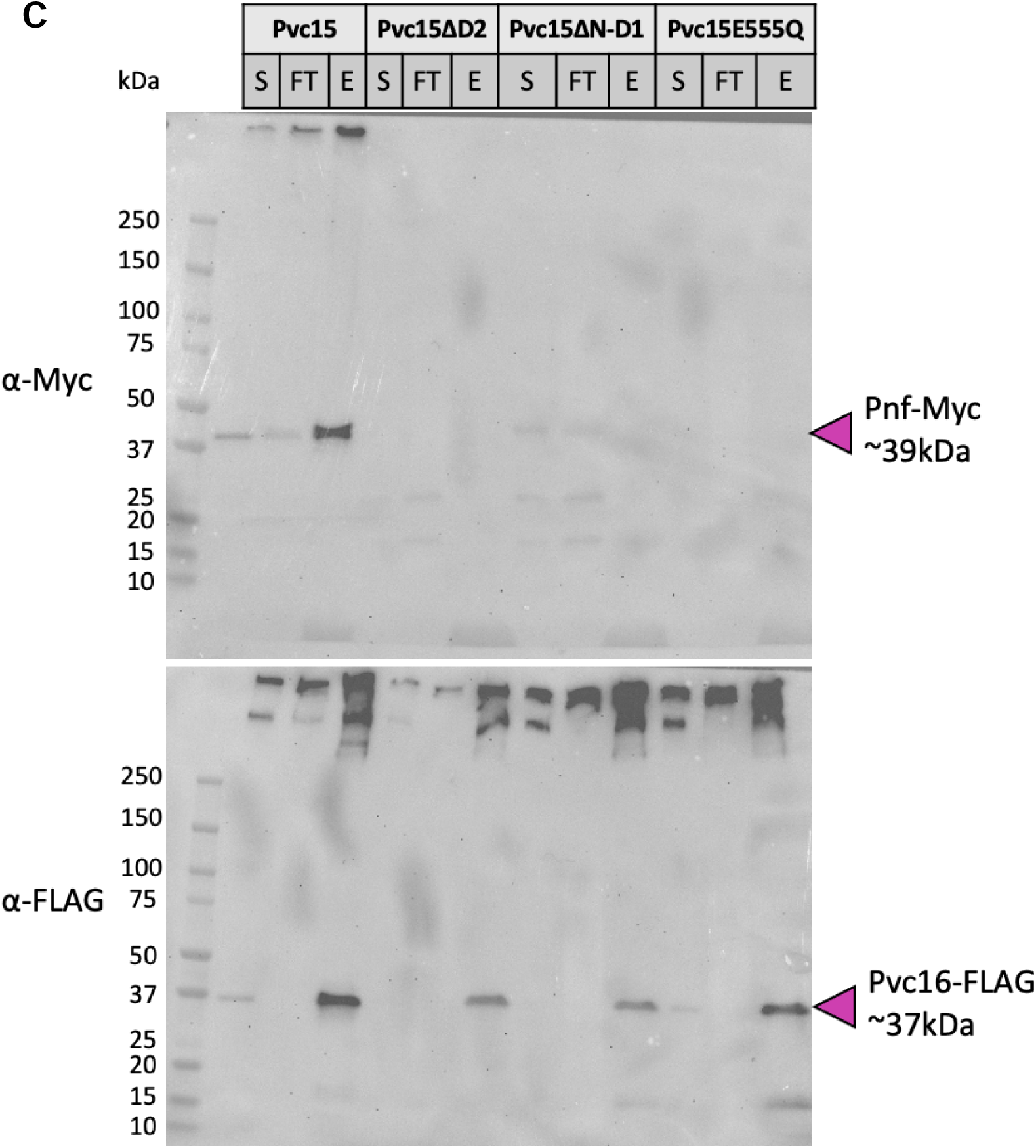
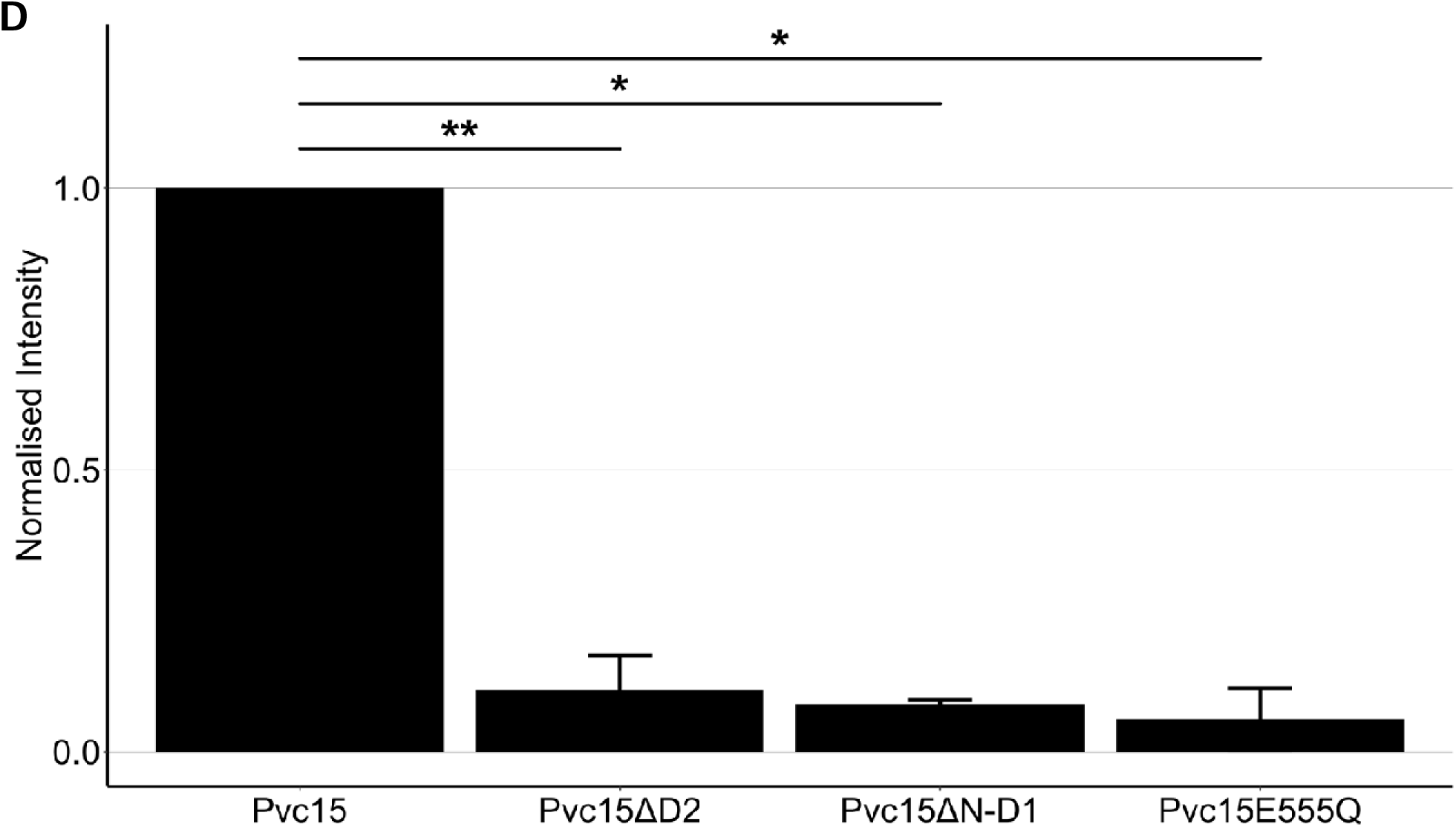
Pvc15 domain D2 stabilises the payload and has PVC*pnf*-dependent ATPase activity for cargo loading. pVTRaDuet, encoding Pvc15 and Pnf downstream of the same promoter, was expressed in *E. coli* and used to measure whether Pvc15 mutations influenced **(A)** Pnf cell lysate abundance, **(B)** ATPase activity, and **(C)** cargo loading. **(A)** Pnf co-expressed with Pvc15 lacking domain D2 had depleted abundance compared to the wild-type (t(4) = -8.66, p-value = 0.0010 ***). The Pvc15 D2 B-box mutant, E555Q, retained a high level of Pnf in lysates compared to when the whole domain was deleted (t(4) = -4.68, p-value = 0.0094 **), implicating D2 as an essential chaperoning domain whilst N-D1 are less efficient chaperones. **(B)** A colorimetric malachite green ATPase assay (Abcam; Cat. No.: ab234055) was done in a 96-well plate to analyse *E. coli* lysates. Only the wild-type Pvc15 co-expressed with Pnf and the PVC*pnf*Δ*pvc15* operon was shown to have significant ATPase activity. Comparing to the Pvc15+Pnf PVC*pnf*Δ*pvc15*^+^ strain: *E. coli*+Pnf: t(4) = -4.59, p-value = 0.010 *; Pvc15ΔD2: t(4) = -4.29, p-value = 0.013 *; Pvc15ΔN-D1: t(4) = -4.08, p-value = 0.015 *; Pvc15E555Q: t(4) = -4.37, p-value = 0.012 *. Mutants were not significantly different from that of the *E. coli*+Pnf negative control, whilst the pVTRaDuet + pBADPVC*pnf*Δ*pvc15* strain was not significantly different from the positive control (t(3) = -1.05, p-value = 0.37). Furthermore, wild-type Pvc15 only retained its level of ATPase activity when co-expressed with the PVC*pnf*Δ*pvc15* operon (with vs without PVC*pnf*Δ*pvc15*: t(4) = -3.18, p-value = 0.033 *). Error bars show **standard error of the mean**; statistical inference was done via a two-sided Student’s T-test; ns = not significant, * = p < 0.05. Data were collected as 3 replicates of separately cultured *E. coli* lysates. **(C)** PVC*pnf* was purified by co-immunoprecipitation with an epitope tagged PVC protein whilst co-expressing Pvc15 mutants. PVC*pnf*Δ*pvc15* strains co-expressing ATPase-incompetent Pvc15 mutants showed a lack of associated Pnf payload compared to the wild-type, indicating that ATPase activity was required for effective cargo loading. S = supernatant, FT = flow-through, E = elution (purified PVC*pnf*). Data is representative of 3 reiterations of PVC*pnf* purifications. **(D)** Quantification of **(C)** as well as 2 other reiterations of PVC*pnf* purifications.

To ascertain whether these mutants still had ATPase activity, a malachite green assay was used to measure a change in OD_650_ and normalise to a standard curve (FIG. S10) to calculate the amount of inorganic phosphate generated from lysates when ATPase substrate is introduced (FIG. 5B). Each of the mutants had activity comparable to the negative control (PVC*pnf*Δ*pvc15* + Pnf; 1.9 mU/mL) whilst wild-type Pvc15 co-expressed with the PVC*pnf*Δ*pvc15* operon and the Pnf payload had much larger ATPase activity (23.0 mU/mL). Notably, this ATPase activity was not replicated if lysates did not co-express the remaining PVC*pnf* operon proteins, suggesting that Pvc15’s ATPase activity, conducted solely by the D2 domain but dependent on intact N-D1 domains, also requires interactions with other PVC*pnf* proteins or mature PVC to function. Altogether, this data suggests that the PVC*pnf*-dependent ATPase activity facilitates translocation of payload into the PVC tube lumen.

Indeed, purified PVC*pnf* lacked co-purification with payload Pnf only if co-expressed with the ATPase-incompetent Pvc15 (FIG. 5C), confirming that ATPase activity is required for translocating payload into the tube lumen of the mature PVC.

### The effect of SP mutations on cargo stability

Jiang and collagues (2022) investigated the effects of point mutations within the signal peptide to identify any single residues which may have altered the association of Pnf with the PVC using alanine scanning and failed to identify any key residues that alter loading of Pnf50-*Renilla* luciferase (Rluc). However, this data does not reveal the nature of Pvc15-specific interactions nor how stability of the native cargoes, such as Pnf, are affected by point mutations since Rluc is stable regardless of the presence or absence of these N-terminal SPs.

It was investigated whether mutations to the SP would confer instability to the cargo with or without Pvc15. To demonstrate each set of point mutations, a pVTRA vector encoding both the native Pnf downstream of double polyhistidine-tagged Pvc15 was constructed and termed pVTRaDuet (Figure S8A). However, α-His Western Blot did not indicate successful expression of Pvc15 at 4 h post-induction or overnight incubation (data not shown). Since the previous pVTRA vector indicated that Pnf is highly unstable at 4 h post-induction in the absence of Pvc15, however, the expression profile of the pVTRaDuet vector (Figure S8B) shows that the stabilising effect of Pvc15 is present despite not being expressed at detectable levels in Western Blot analysis.

Interestingly, Pnf co-expressed with Pvc15 for 20 h post-induction results in loss of detectable cargo; however, cargo levels can be reconstituted when also co-expressed with the rest of the PVC*pnf* operon (Figure S8B; lanes 4-6 compared to lanes 10-12); there was no difference in Pnf-Myc band intensity at 4 h post-induction of the pVTRaDuet vector with or without the co-expression of the PVC*pnf*Δ*pvc15* operon. This may suggest additional interactions mediate cargo stability after overnight PVC induction, or that translocation into the PVC tube lumen is necessary for retaining payload abundance in the cytosol after long incubations.

## Conclusion

This work showed that each of Pvc15’s domains are vital for associating SP-associated cargo proteins with the PVC, and that SPs are ubiquitous in other homologous eCIS putative cargoes such as the Afp in *Serratia* and *Yersinia*. Despite their similar role in PVC maturation, these SPs act differentially with the Pvc15 ATPase and chaperone; their groups of homology between similar strains and operons in *Photorhabdus* may give clues as to whether certain conserved motifs within SPs have similar roles or protein-protein interactions.

The N-domain was found to have homology to DNA-binding proteins, but what role such activity has in Pvc15’s role should be investigated further. The D2 domain, in particular, plays significant roles in the ATPase activity, SP stabilisation, and PVC loading, whilst the D1 domain may be involved in stabilising the Pvc15 hexamer. In this work, Pvc15 ATPase activity was demonstrated to mediate the loading of SP-associated payload into the PVC. Understanding the functional roles of the SP, Pvc15, and interactions with other PVC components will enable researchers to facilitate the potential future use of these devices for eukaryotic inter-membrane trafficking of bioactive therapeutic proteins.

## Materials and Methods

### Plasmid constructs used in this work

The pVTRA vector (Pérez-Martin and de Lorenzo, 1996) was used for cargo expression by the addition of 0.1-1.0mM IPTG. The pBAD-PVC*pnf* vector was constructed by synthesis of the PVC*pnf* operon encoding FLAG (DYKDDDK) to the C-terminus of Pvc16 and can be induced with 0.2% arabinose (Guzman *et al*., 1995). An epitope tag on one of the PVC*pnf* proteins was also used for immunoprecipitation of mature PVC*pnf*. The pVTRaDuet vector, which encodes Pvc15 and Pnf under the control of the same promoter, was generated by PCR of Pvc15 with overhangs encoding hexahistidine tags and the relevant restriction enzyme sites into pVTRA: KpnI and BamHI using the primers (Forward: 5’-GGTACCTTGACCACAATGACTTAGTCTGAGTAAAAAATATGCACCACCACCACCACCACAATATATCGCCTGTTTTTTATGATTCATTG-3’; Reverse: 5’-GGATCCTCCTCAATGATGATGATGATGATGAAATGTTAATCGTCCGACTTTAGC-3’).

### Analysing the dbeCIS effector dataset

Using R within RStudio (R Core Team, 2020; RStudio Team, 2020), regular expressions (via the *grepl()* function) were used to count the occurrences of chosen keywords that described the most common predicted function entries in the dataset. Of the entries that did not include the set of keywords, the phrases ‘t\\dss’, ‘toxin’, or ‘effector’ were used to infer the ORF as a toxin, designated as ‘Other Toxin’, or otherwise assigned as ‘Other’. If different functions were predicted for a single ORF, the one that appeared most commonly was preferred.

Function counts were normalised by the number of strains within each species as well as the number of operons in each genome. For lineage Ib, the counts were also normalised by the number of species in the dataset. Thus, one can infer the frequency of each function for the expected number of ORFs encoded within 5kb downstream of any given eCIS operon from any given strain in that genus.

### Protein expression

Transformed BL21(DE3) *E. coli* (New England Biolabs) (NiCo21) were incubated overnight in a starter culture of 10mL at 180rpm and 37 °C with the appropriate antibiotic(s); each biological replicate was grown into a separate starter culture to minimise inter-sample interference. 5mL of this culture was diluted 1:100 and grown again for 3-4 h to OD_600_ = 1.0 at which point 0.2% arabinose was used to induce expression of the pBAD-PVC*pnf* vector and 1mM IPTG was used to induce the pVTRA vector encoding the cargo. Cells were then incubated at 200rpm and 25 °C.

For measuring cargo stability, aliquots were taken 3 h post-induction; for measuring degradation of the cargo, kanamycin was added at a final concentration of 20μg/mL and aliquots were taken after 4 h. For PVC purification, cells were incubated overnight after induction then centrifuged at 3,000g for 20 minutes at 4 °C. Cells were then lysed using 10mL of lysis buffer per gram of pellet, prepared as follows:

- 200μg/mL lysozyme
- 0.5% Triton X-100
- 1 tablet complete EDTA-free protease inhibitor per 50mL
- 70μL per 100mL DNase I

After resuspension, cells were homogenized using a cell homogenizer at 30kpsi (One Shot Model; Constant Systems LTD).

For making aliquots at each stage of the purification process, 25μL of sample was taken and added to 25μL 2X LDS sample buffer with 50mM DTT and boiled for 20 minutes at 95 °C. The homogenized sample was then spun at 12,000g for 35 minutes at 4 °C and the supernatant was filtered through a syringe filter with 0.45μm pore size.

PVC syringes were purified via co-immunoprecipitation of one of the PVC proteins with a gravity flow column (Bio-Rad Econo-column^®^; cat. no.: 7371512), used according to the manufacturer’s instructions.

### SDS-PAGE, Western Blot analysis, and quantification

Sodium-Dodecyl-Sulphate (SDS) Polyacrylamide Gel Electrophoresis was done in precast gels (Bio-Rad TGX Mini-Protean 4-15% gels with Dual Colour Standard, cat. no.: 4561084DC) which were used as per the manufacturer’s instructions. Runs were conducted in a Mini-Protean gel tank using PAGE buffer and run at 150V for 45 minutes.

After gel running, the extracted gel was loaded onto a PVDF membrane using a Biorad Transblot Turbo electroblotter for 7 minutes. The Thermo Pierce Fast Western Blot kit (cat. no.: 35055) was used according to the manufacturer’s instructions. Rabbit anti-FLAG (DYKDDDDK, Cell Signaling, 14793) or mouse anti-Myc (Cell Signaling, 2276) monoclonal antibodies were used and visualised using the horseradish peroxidase (HRP)-bound secondary antibody provided in the Pierce kit after adding luminol and peroxide buffer. Bands were visualised using the chemiluminescence protocol in a G:BOX -F3 (Syngene) and using 2x2 binning for more than 300 seconds of exposure.

For quantification of band intensities, samples were normalised based on their OD_600_ for cell lysates or protein concentration for purifications, as determined using the BSA protocol on a spectronanophotometer. In addition, for a given band, the mean background value of the blot was subtracted and a corresponding loading control was used for that replicate either in the form of a fixed point band which appeared in all samples, such as the FLAG-tagged Pvc16 cap protein, or the mean of all points in the corresponding lane in the Coomassie blue stain calculated using FIJI (ImageJ). After dividing the mean grey values of each band by its corresponding loading control mean grey value, test bands were relativised to their control samples; statistical analysis was done using a Student’s T-test.

### Resazurin assay for cell respiration

Raji cells incubated at 5% CO_2_ and 37 °C for 2 days were diluted to 8 x 10^5^ cells/mL (determined by first conducting a calibration curve experiment to optimise the starting cell number) and seeded at 100μL in quadruplicate in a black, clear-bottomed 96-well plate. Background fluorescence from the media was subtracted from the mean fluorescence of the 9 media-only wells at each time point. The plate was incubated for 1 h at 5% CO_2_ at 37 °C and PBS or a sample of 10μg of PVC was added (measured using a nanospectrophotometer) and incubated for a further 4 h. Finally, resazurin (12.5mg/mL stock) was added to each well in quick succession (∼5μM final concentration). The plate was incubated for a further 6 h and fluorescence was measured: excitation = 530-570nm, emission = 580-620nm.

### ATPase activity

The colorimetric ATPase assay kit (Abcam, cat. no.: ab234055) was used with cell lysates, as per the manufacturer’s instructions. 15mL of *E. coli* NiCo21 cultures grown to OD600 = 1.0 were induced with 0.2% arabinose and 0.5mM IPTG to co-express PVC*pnf*Δ*pvc15* and the LS-associated payload with or without Pvc15 for 16 h then processed according to the manufacturer’s instructions.

## Acknowledgements

This work was supported by the Biotechnology and Biological Sciences Research Council (BBSRC) as part of the Midlands Integrative Biosciences Training Partnership (MIBTP); grant number: BB/M01116X/1. Thanks go to Dr. John James for assisting with experimental design and providing certain materials and cell lines and to the collaboration with Nanosyrinx Ltd.

## Supplementary Material

**Figure S1.**

PVC signal peptide homology is more aptly described in distinct groups than all-vs-all alignments.

**Figure S2.**

AlphaFold2.0 predictions of native PVC cargo effectors display signal peptides as disordered polypeptides.

**Figure S3.**

Tertiary overlay of Pvc15 homologues indicates high structural conservation, particularly in the D1 and D2 domains.

**Figure S4.**

D2 Walker A P-loop likely coordinates phosphate in its ‘LRLR’ nest.

**Figure S5.**

A complete prediction of a Pvc15 hexamer reveals an open right-handed spiral staircase.

**Figure S6.**

Pnf expression calibration curve indicates optimal payload expression at 3-4 h post-induction.

**Figure S7.**

Raji B cell resazurin assay calibration curve of starting cell number.

**Figure S8.**

pVTRaDuet vector for simultaneous Pnf-Myc and Pvc15 expression may indicate additional PVC core interactions.

**Figure S9.**

Induction of the PVC*pnf* operon does not affect the ability of wild-type Pvc15 to rescue Pnf abundance.

**Figure S10.**

Standard curve for conversion of OD_650_ to nmol phosphate at t=0.

