## supplementary information for "The Pvc15 D2-Pnf SP Interaction Mediates AAA ATPase Activity, Payload Stability, and Translocation into the PVC Tube Lumen"

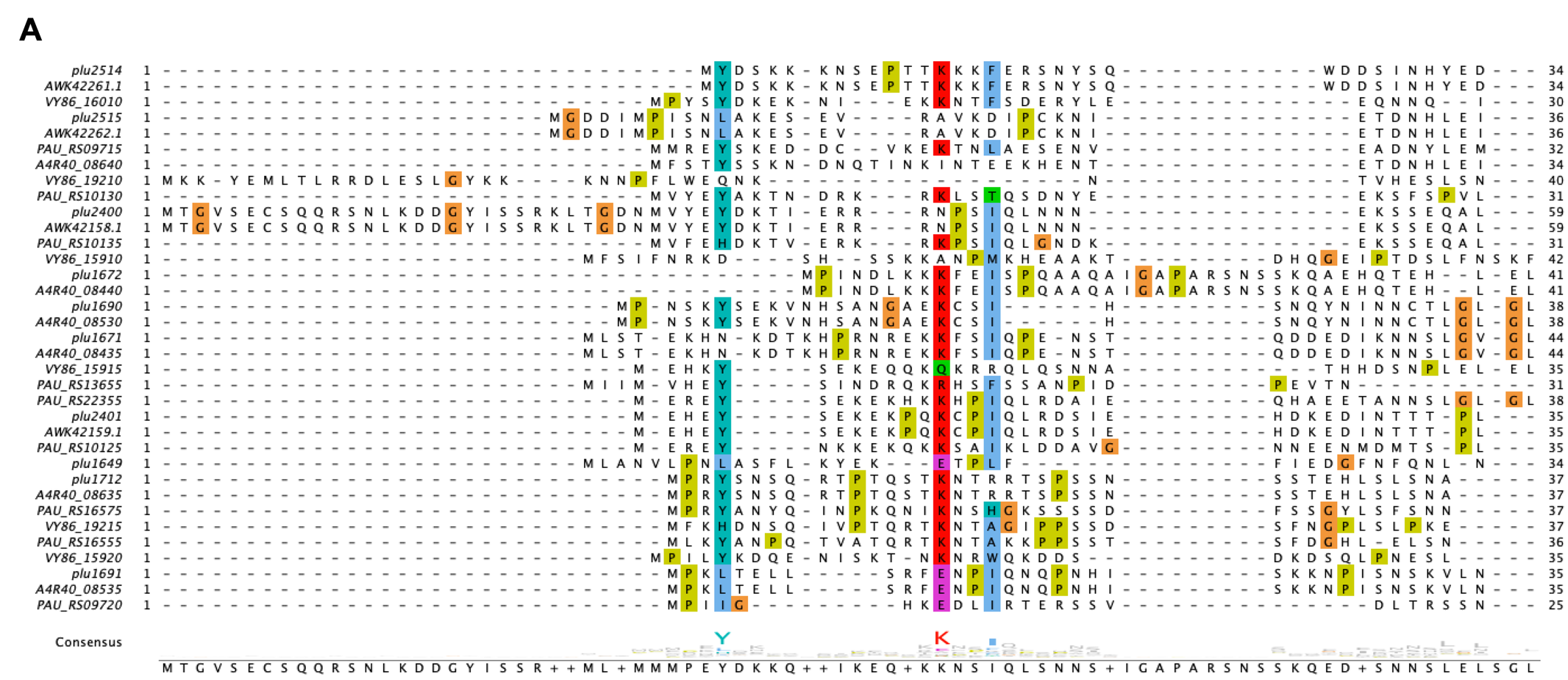

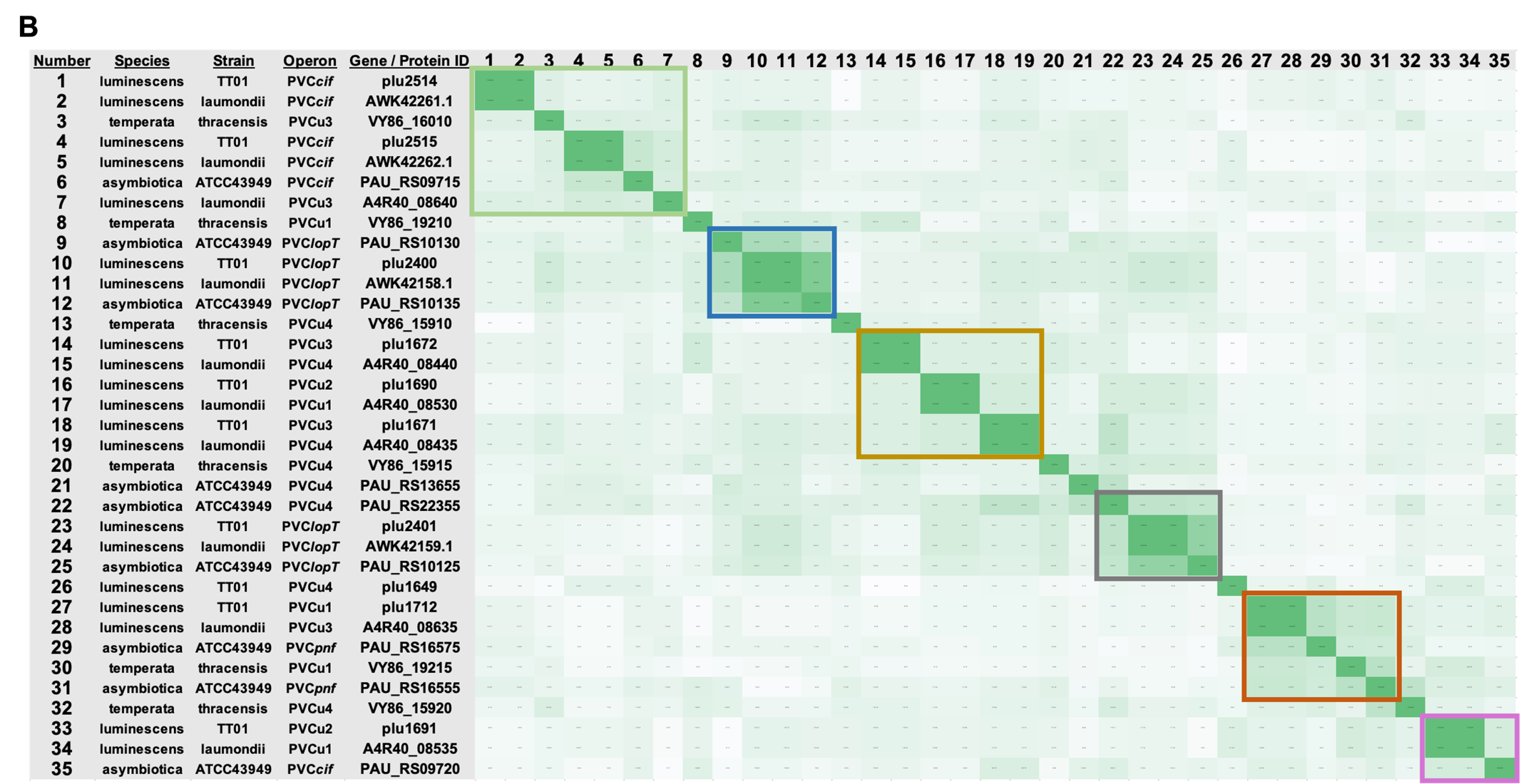

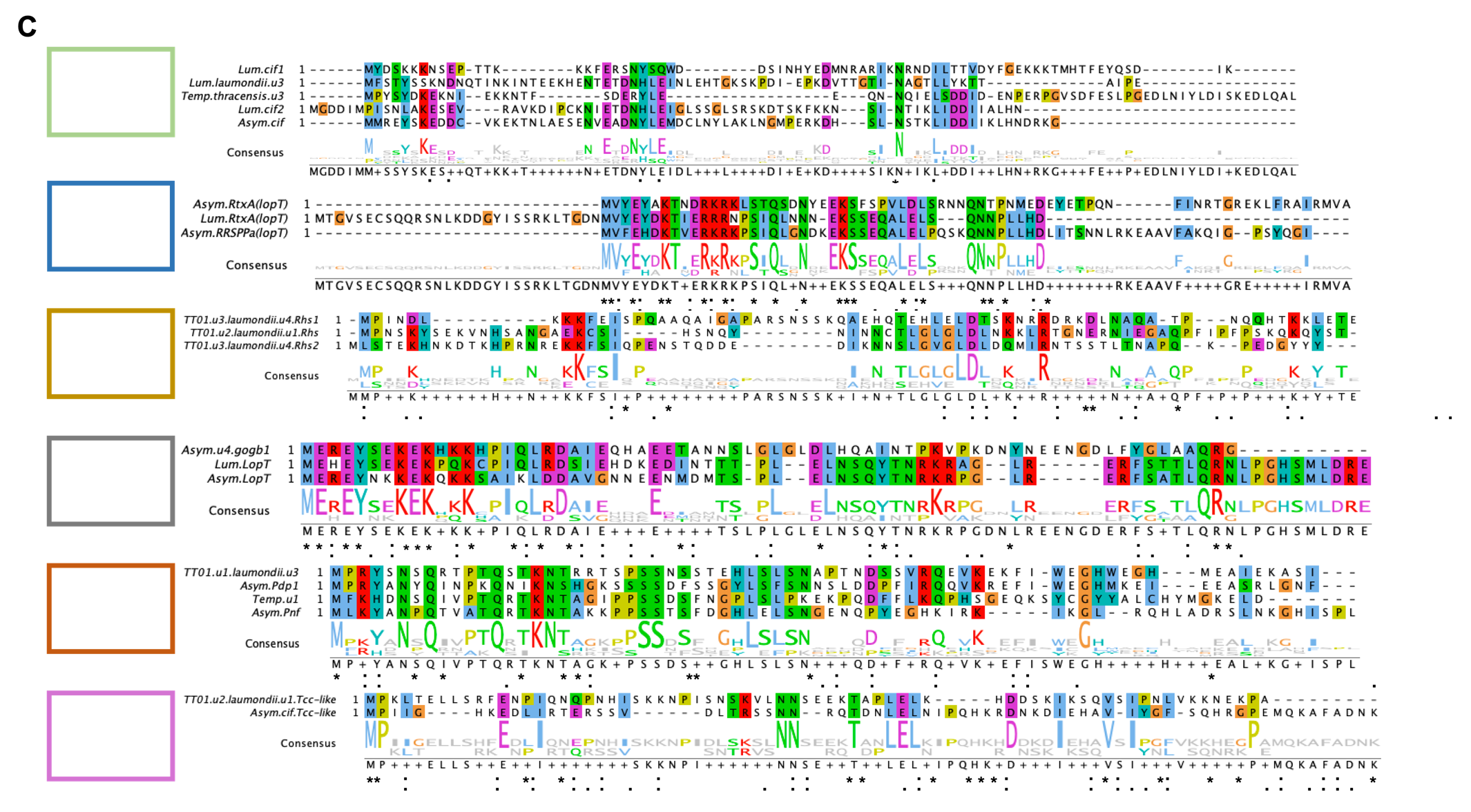

FIG. S1. PVC signal peptide homology is more aptly described in distinct groups than all-vs-all alignments.

**(A)** All-vs-all local alignment in Clustal Omega of the signal peptides (SPs) of PVC-downstream ORFs, selected from the dbeCIS dataset used in FIG 1, only vaguely demonstrates the existence of rarely conserved residues, as described by Jiang and colleagues (2022). **(B)** Construction of a percentage identity matrix and colouring by scores in the form of a heatmap displays more close groupings of homology that may better describe the nature of these SPs. **(C)** Local alignment of these groupings indicates conserved *motifs* within SPs belonging to a given group which may be better descriptors of homology than conserved *residues*.

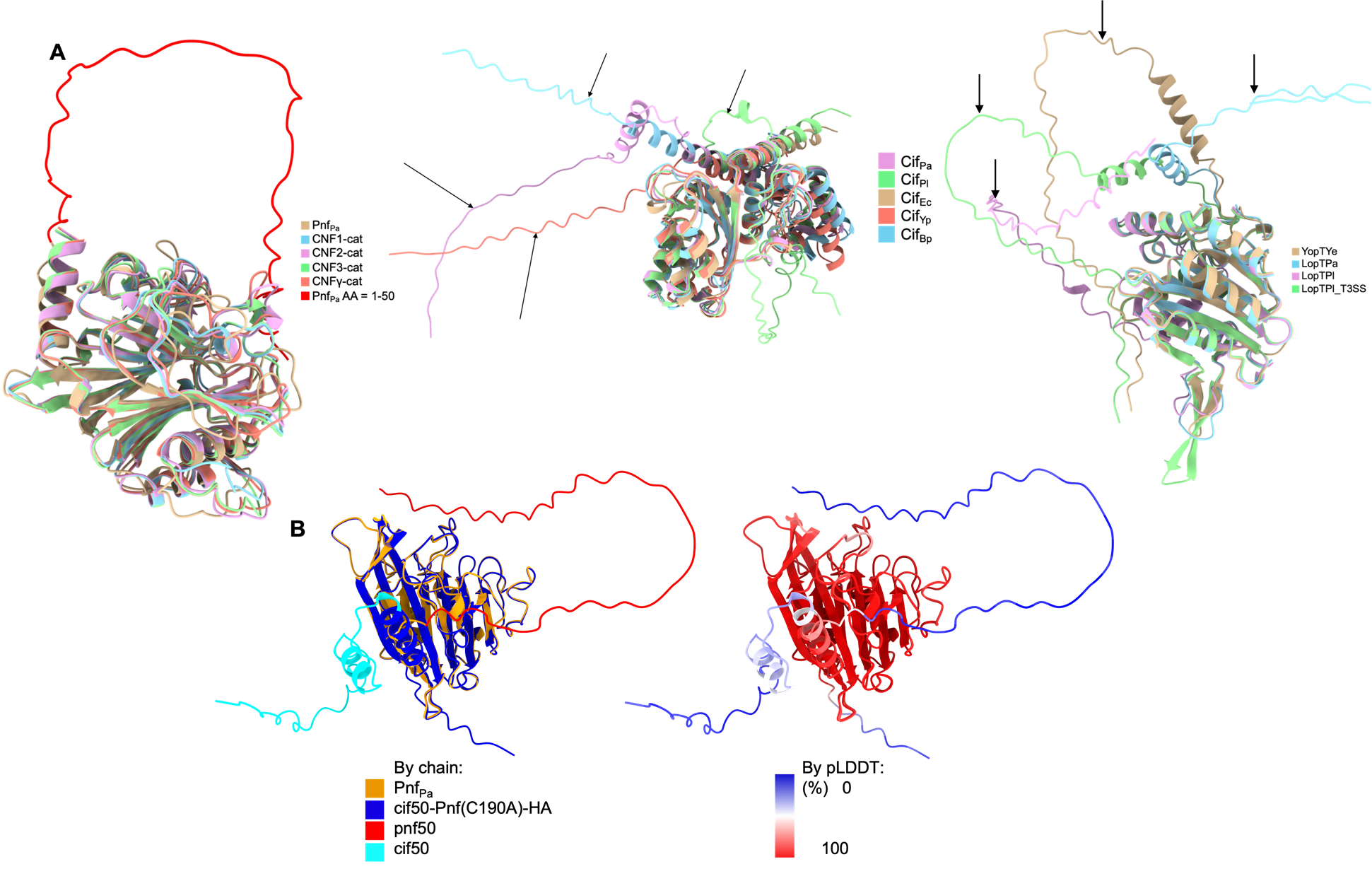

FIG. S2. AlphaFold2.0 predictions of native PVC cargo effectors display leader sequences as disordered polypeptides.

The AlphaFold2.0 confidence value, predicted local distance difference test (pLDDT), can be used to infer the presence of leader sequences; N-terminal regions with particularly lower pLDDT values, typically less than 40%. **(A)** Native PVC effectors Pnf, Cif, and LopT are aligned with the structures of their homologues and show distinct disordered leader sequences: i) the catalytic domains of cytotoxic necrosis factor (CNF-cat) from *E. coli* (CNF1-3) and *Y. pseudotuberculosis* (CNFγ); ii) Cif proteins from *P. asymbiotica* ATCC43949 (Cif_Pa_), *P. luminescens* TT01 (Cif_Pl_), *E. coli* (Cif_Ec_), *Y. pseudotuberculosis* (Cif_Yp_), and *B. pseudomallei* (Cif_Bp_); iii) YopT proteins from *Y. enterocolitica* (YopT_Ye_), *P. asymbiotica* (LopT_Pa_), and *P. luminescens* loci at either the PVC operon (LopT_Pl_) or encoded at the type 3 secretion system operon (LopT_Pl__T3SS). **(B)** A prediction of the epitope tagged Pnf toxoid used for experiments fused with either its native pnf50 leader (i.e., the first 50 amino acids of Pnf) or the Cif leader, cif50, indicates that both leaders display a low confidence prediction of amino acids which are separated in space from that of the more globular protein structure.

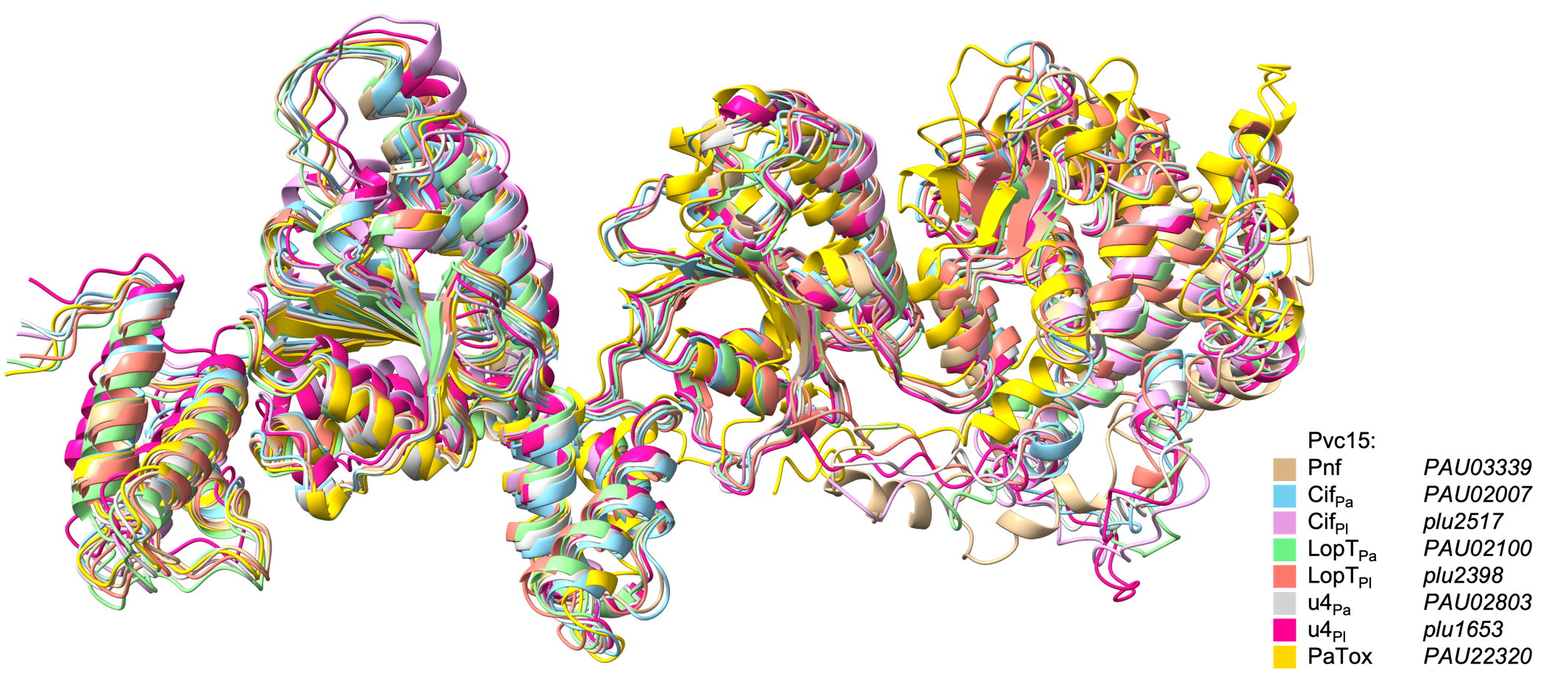

FIG. S3. Tertiary overlay of Pvc15 homologues indicates high structural conservation, particularly in the D1 and D2 domains.

Amino acid sequences for each of the Pvc15 units in *Photorhabdus asymbiotica* ATCC43949 (Pa) or *P. luminescens* TT01 (Pl) were predicted using AlphaFold2. The highest-scoring model was then overlaid with homologues using Matchmaker within the ChimeraX suite; the PVC*pnf* Pvc15 (PAU03339) was used as a reference. AlphaFold2.0 predictions of each of the Pvc15 homologues showed a vastly similar overall structure, though N-domains represented the lowest R.M.S.D.

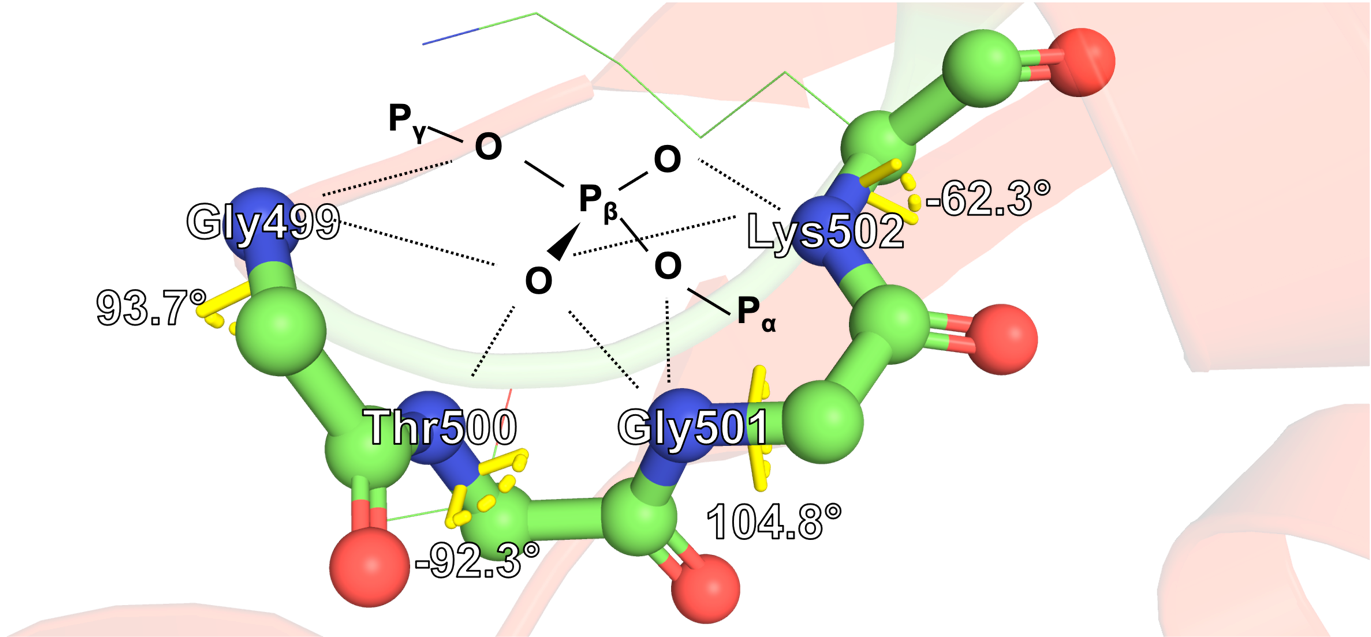

FIG S4. D2 Walker A P-loop likely coordinates phosphate in its 'LRLR' nest.

Adapted from Watson and Milner-White (2002). The Walker A motif “LRLR” nest with proposed coordination of phosphate as documented for the P-loop of Ras. Phi bond angles are also shown; carbon atoms are shown in green, nitrogen in blue, and oxygen in red; Amber forcefield-relaxed side chains are depicted as thin sticks.

**A**

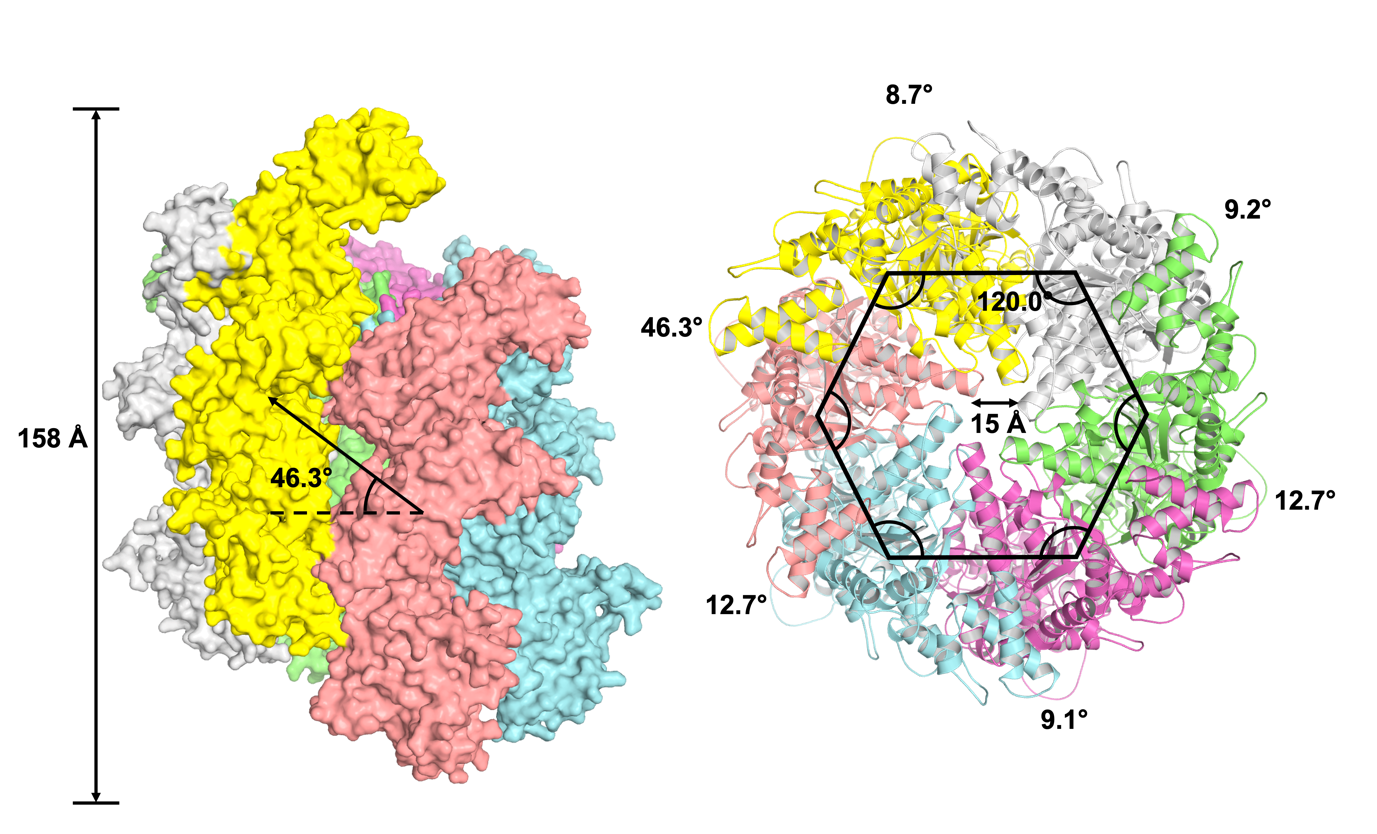

**B**

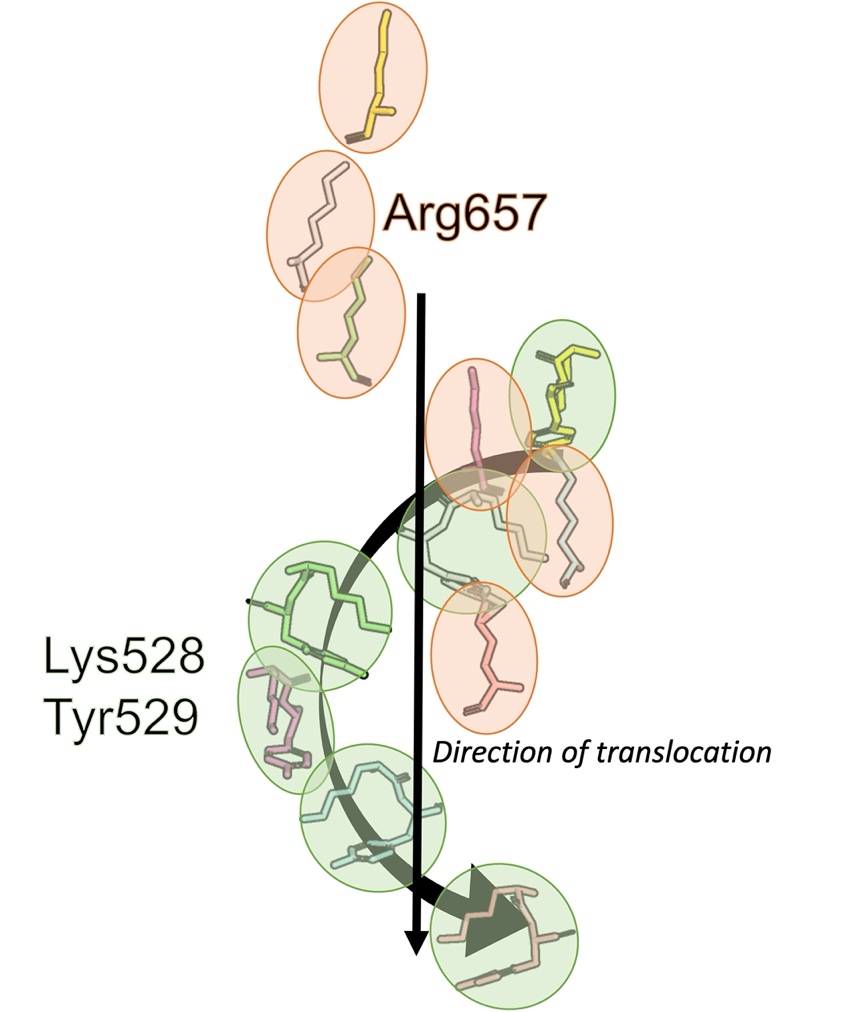

**C**

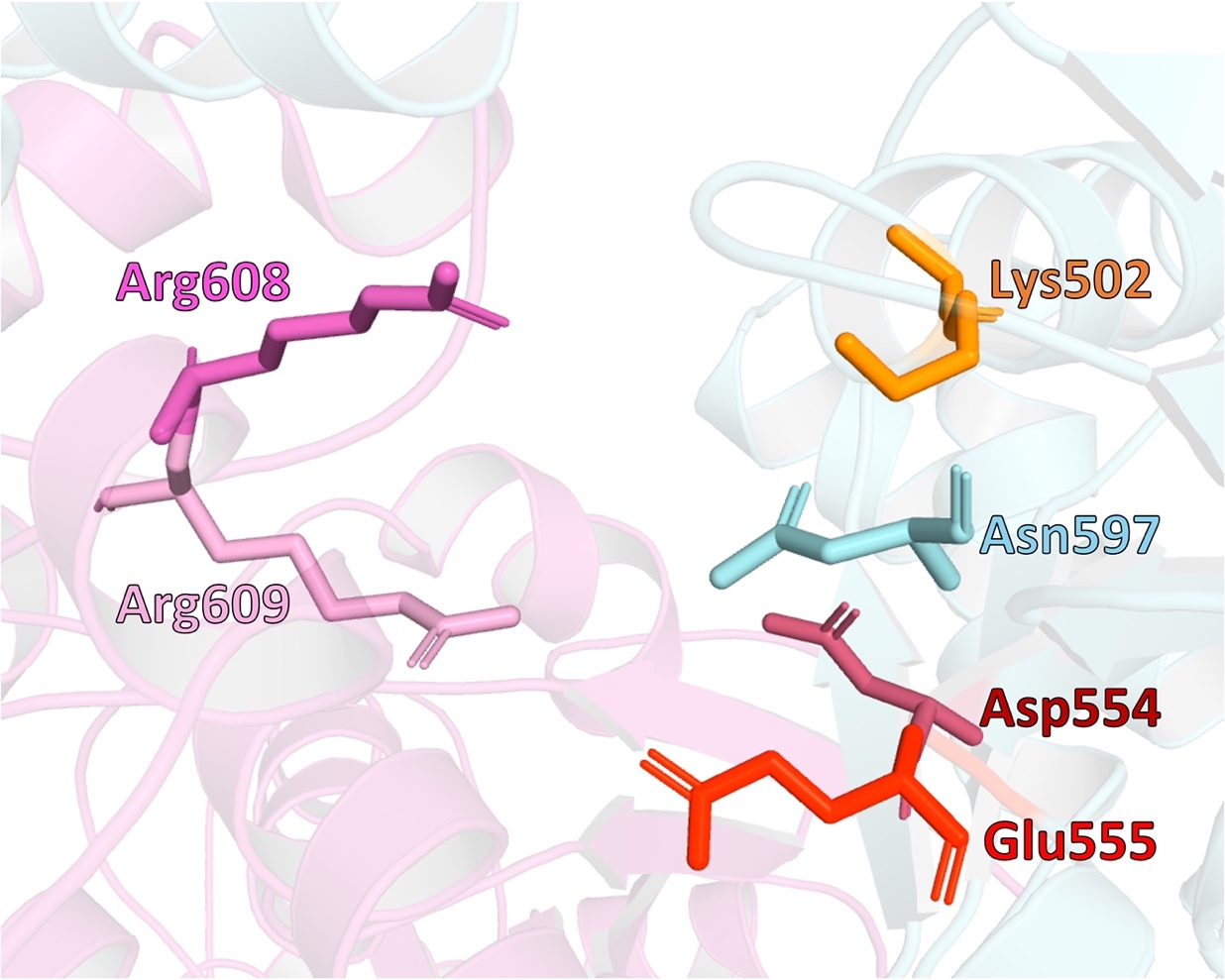

FIG S5. A complete prediction of a Pvc15 hexamer reveals an open right-handed spiral staircase.

Using MoLPC, the hexameric Pvc15 structure could be predicted with high confidence by optimising the predicted local distance difference test (pLDDT) at interfacing residues (pLDDT = 61) indicated by the mpDockQ score of 0.233. **(A)** Pvc15x6 assumes an open, right-handed spiral staircase conformation when not bound to substrate and ATP. The C-terminal D2 domain (top of the left-hand structure) forms a pore with a size of roughly 15Å, similarly to other classical AAA^+^ ATPases. **(B)** Pore loops 1 and 2 are situated such that both can interface with substrate from different sides whilst each subunit sequentially provides an interaction that would translocate the substrate upon ATP hydrolysis. **(C)** R-fingers R608 and R609 are located within 10Å of the predicted ATP-binding site in the hexamer model, acting *in trans* to coordinate ATP during hydrolysis and communicating biochemical events to the adjacent subunit.

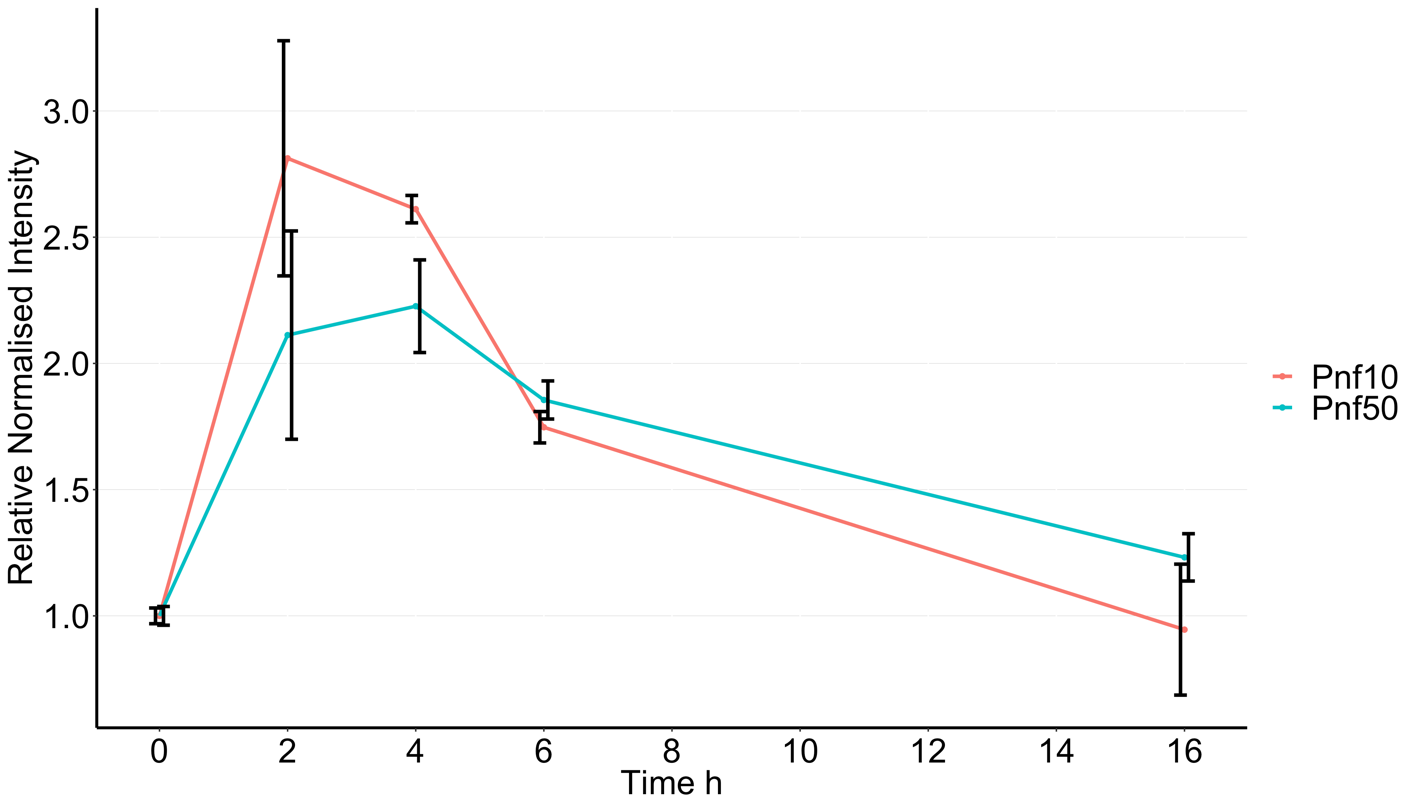

FIG S6. Pnf expression calibration curve indicates optimal payload expression at 3-4 hours post-induction. Western Blot band intensities, normalised by the sum of Coomassie blue stains, were measured after induction of the Pnf payload from the pVTRa vector as well as the PVC*pnf* operon from the pBAD vector with 1mM IPTG and 0.2% arabinose, respectively. Subsequent experiments to measure Pnf abundance and stability were done 3.5 hours post-induction.

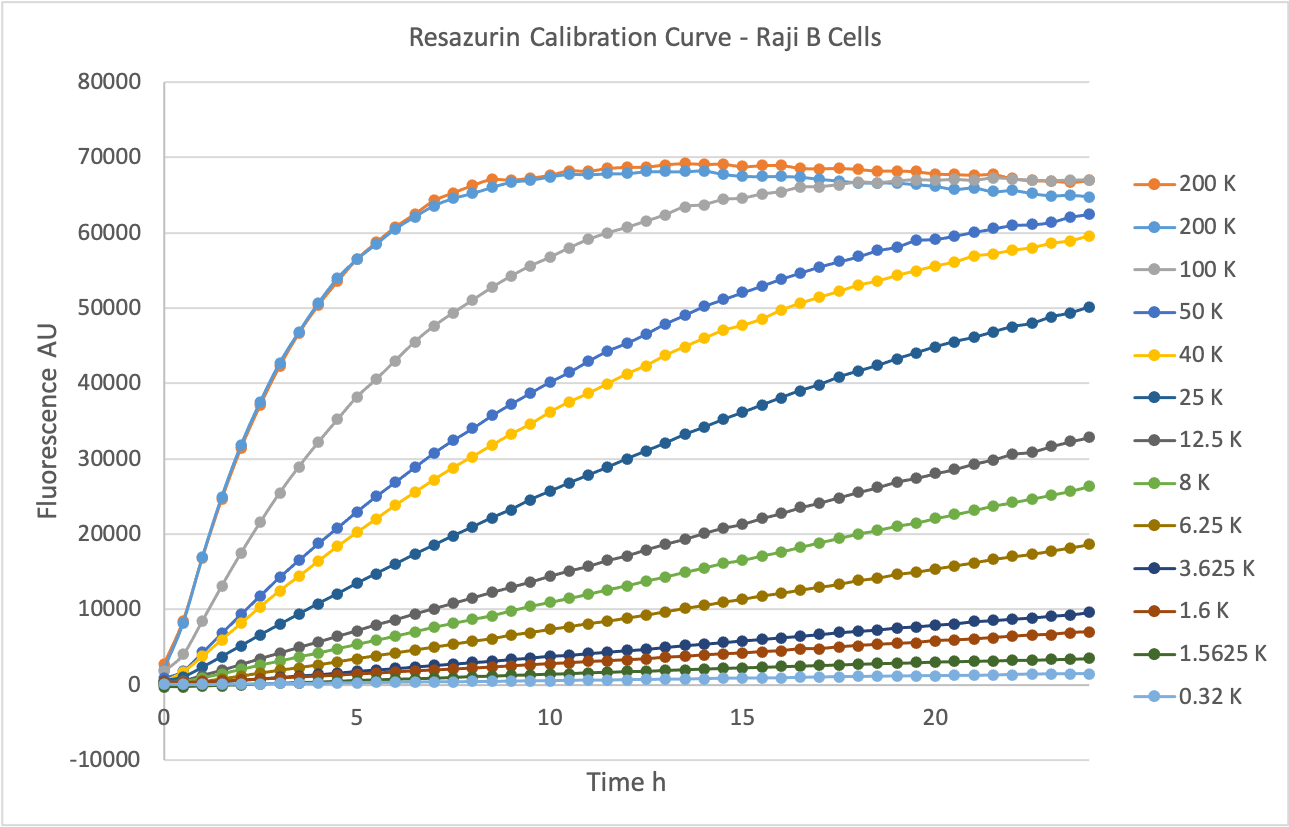

FIG. S7. Raji B cell resazurin assay calibration curve of starting cell number.

The appropriate number of starting Raji cells in a 96-well plate was assessed over a 24 hour time course by measuring the fluorescence of the reduced form: resorufin. Around 70-80K starting cells leads to a plateau in the data at around the end of the time course (100μL per well; 7-8 x 10^5^ cells/mL).

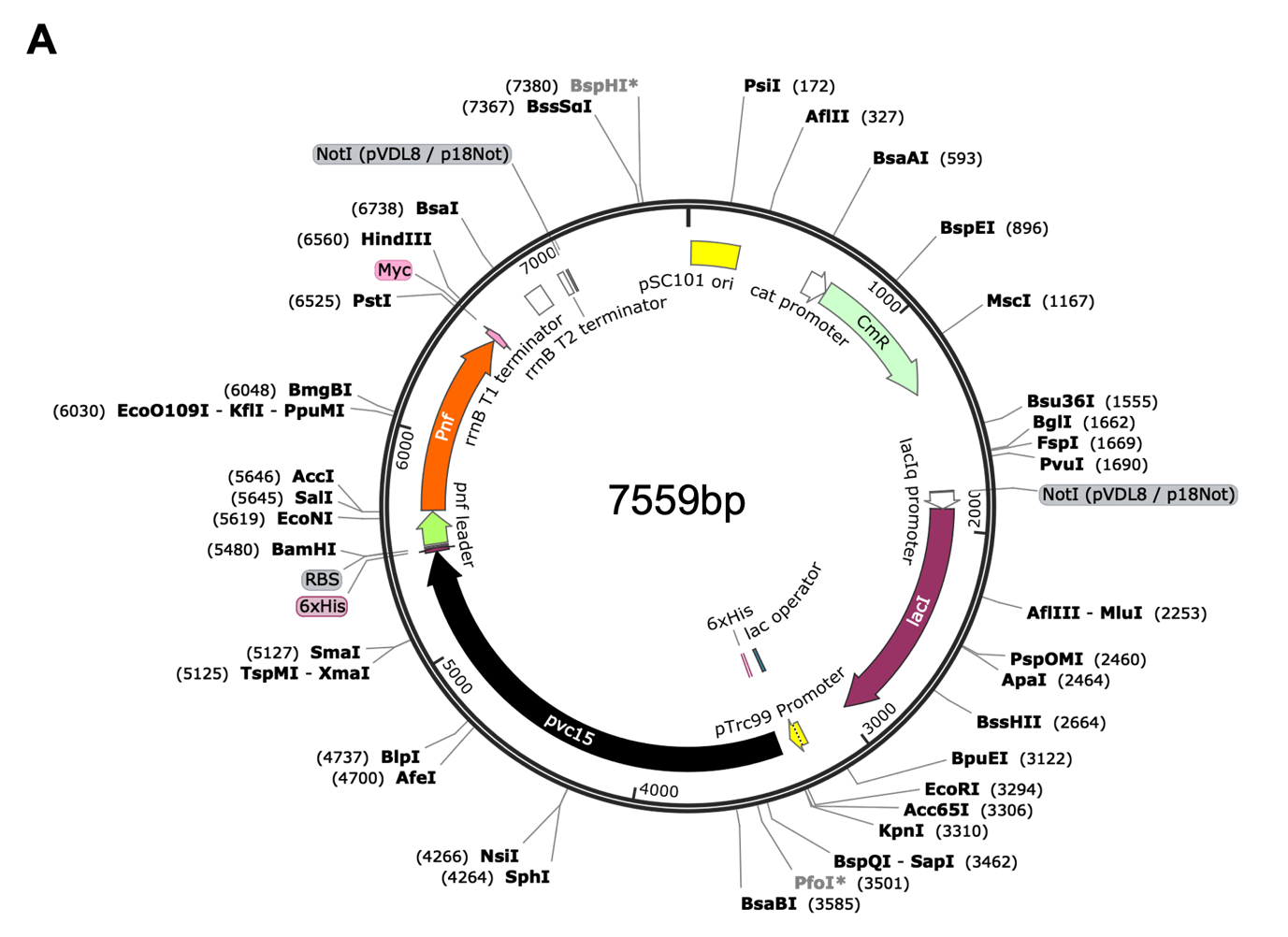

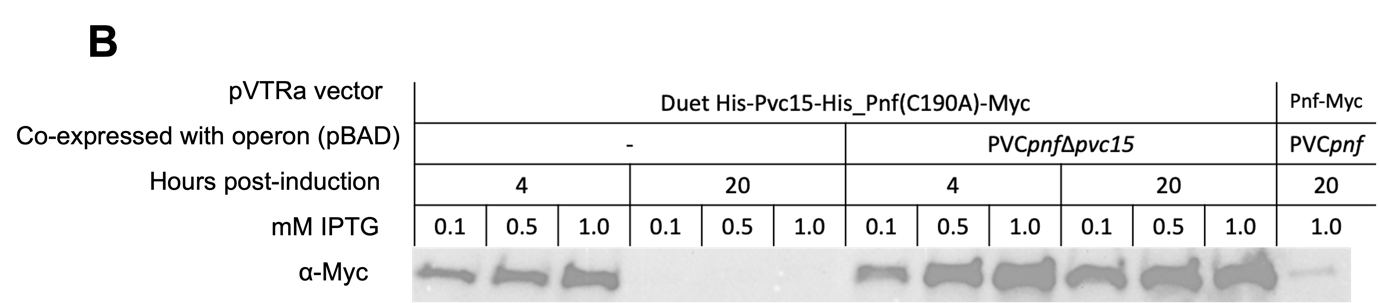

FIG. S8. pVTRaDuet vector for simultaneous Pnf-Myc and Pvc15 expression may indicate additional PVC core interactions.

The pVTRaDuet vector was constructed using the KpnI + BamHI sites within the original pVTRa Pnf(C190A)-Myc vector; sequencing revealed an accurate sequence of the native N- and C-terminal 6xHis tagged Pvc15*_pnf_* as well as the C-terminal Myc tagged Pnf toxoid. **(A)** Labelled sequence map of the pVTRaDuet expression vector with chloramphenicol resistance sites (CatR) and LacI encoded upstream of the lacO *cis* repressor element within the pTrc99-type promoter. Restriction sites KpnI, BamHI, SalI, PstI, and HindIII were used to construct the vector from the original pVTRa vector. **(B)** Whilst the pVTRaDuet vector fails to achieve detectable levels of Pvc15 at 4 or 20 hours post-induction with IPTG, Pnf stability at 4 hours is indicative of its low levels of stabilising activity, as seen in previous figures. The loss of Pnf at 20 hours post-induction when induced on its own is not observed if co-expressed with the arabinose-induced pBAD PVC*pnf*Δ*pvc15* expression vector; this may indicate a separate set of interactions resulting in the stabilisation of the Pnf cargo as a result of other structural components of the PVC*pnf* operon. Notably, Pnf expression appears to be increased when expressed via the pBAD-pVTRaDuet system compared to Pvc15 expression in cis with the pBADPVC*pnf* operon (lane 12 and 13).

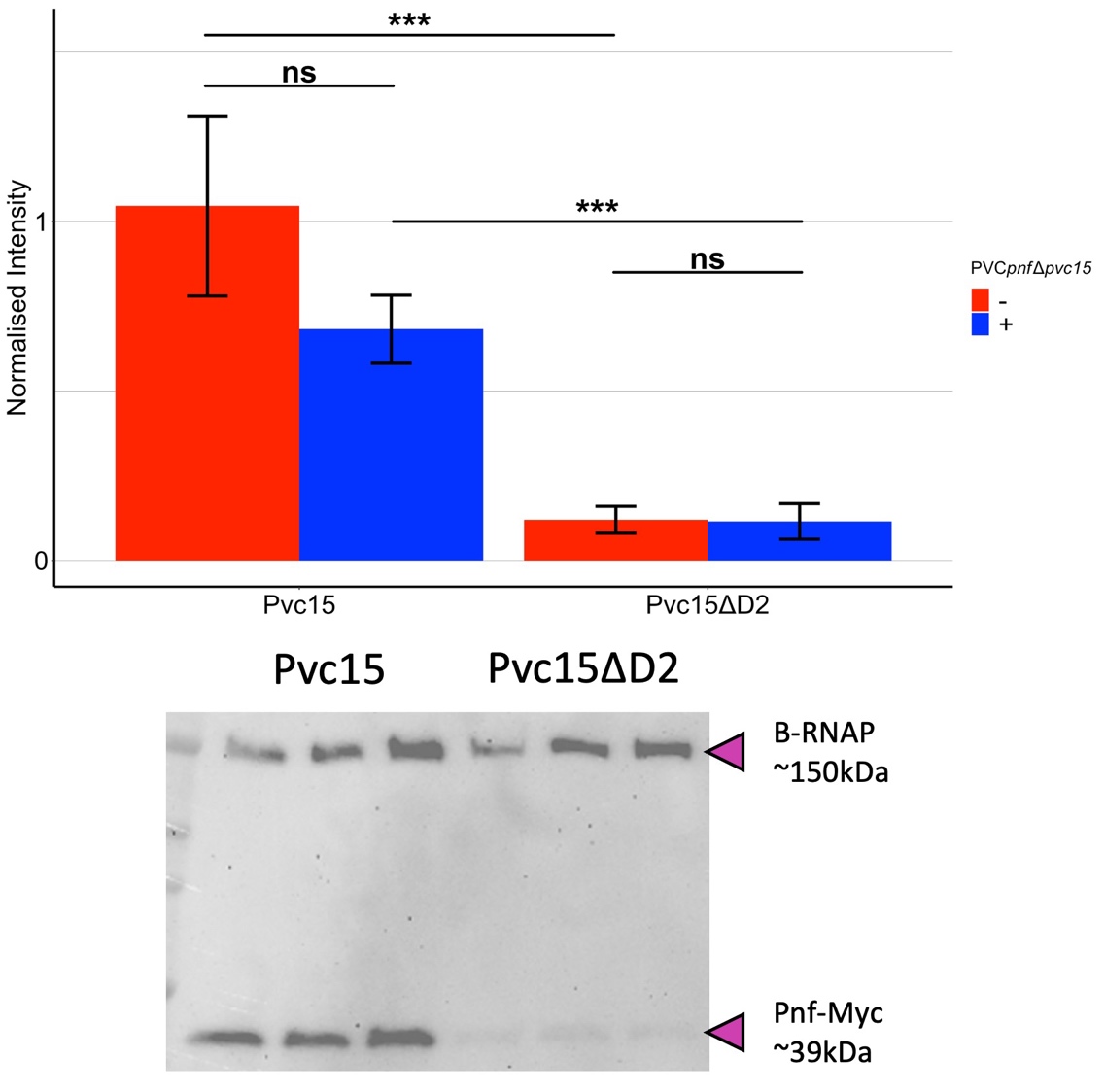

FIG S9. Induction of the PVC*pnf* operon does not affect the ability of wild-type Pvc15 to rescue Pnf abundance.

Lysates from cells expressing Pnf and Pvc15 via the pVTRaDuet vector indicated that the Pvc15 mutant lacking the D2 domain reduced Pnf abundance regardless of whether the cells were co-expressing pBADPVC*pnf*Δ*pvc15* (wild-type and mutants) (without PVC*pnf*Δ*pvc15*: t(4) = -5.97, p-value = 0.0040***; with PVC*pnf*Δ*pvc15*: t(4) = -8.66, p-value = 0.0010***).

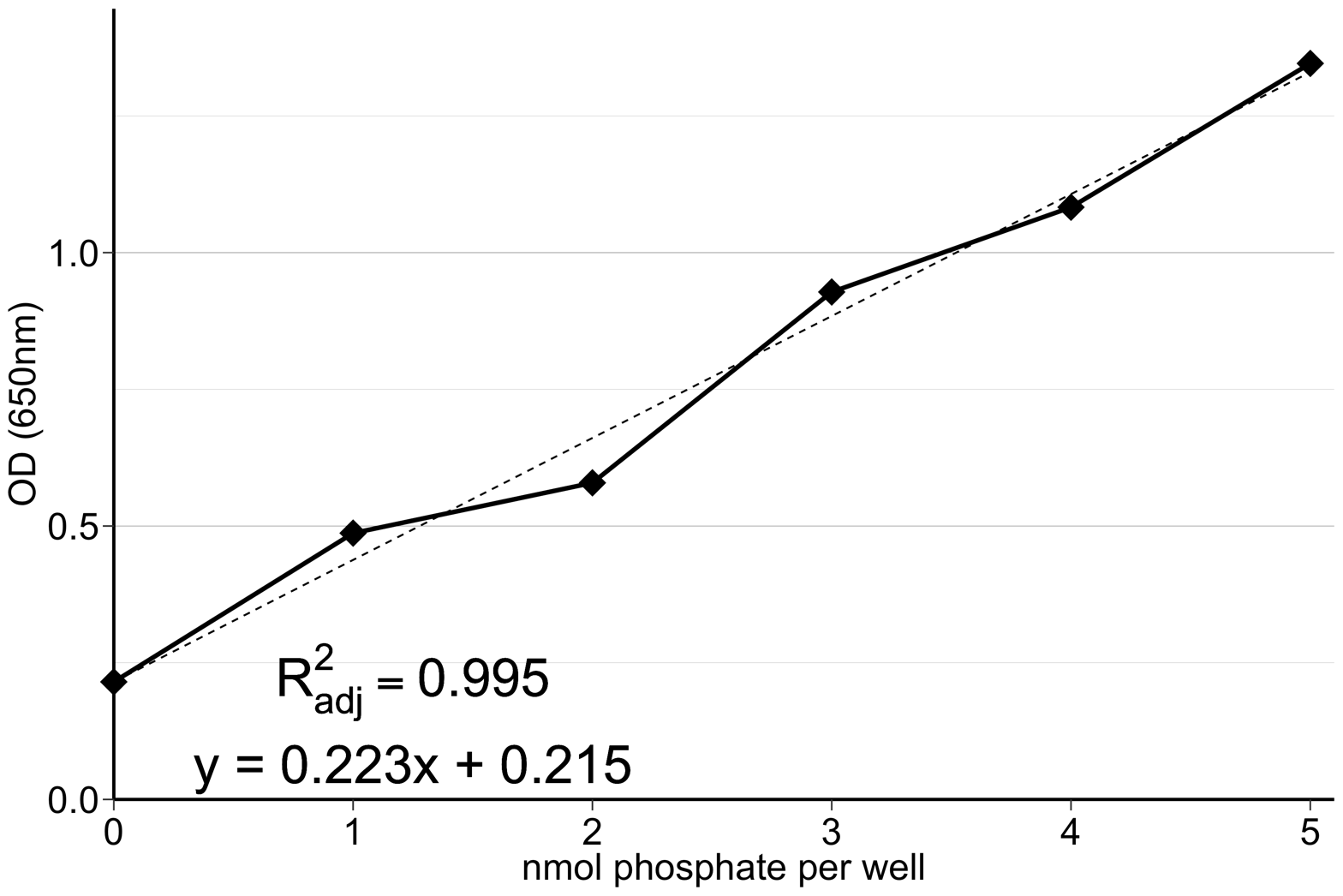

FIG S10. Standard curve for conversion of OD_650_ to nmol phosphate at t=0.

Optical density at 650nm was used to assess how much phosphate was evolved over the 30 minute time course as a measure of the Pvc15 ATPase activity. Change in nmol phosphate could be converted to ATPase activity (U/mL) by dividing by the volume of lysate added to wells and time length of the experiment.
